# Inflammatory arthritis disrupts ocular immune privilege by compromising blood-retinal barrier integrity and promoting uveitogenic T cell recruitment

**DOI:** 10.64898/2026.07.13.738174

**Authors:** Simon Eastham, Amy Ward, Chloe Moscrop, David G. Hill, Aisling Morrin, Sandra Dimonte, Alison Young, Simon A. Jones, Jian Liu, Andrew D. Dick, David A. Copland, Lindsay B. Nicholson, Gareth W. Jones

**Author notes:** **Corresponding author:** Dr Gareth W Jones, School of Cellular & Molecular Medicine, University of Bristol, Bristol, UK. SE and AW contributed equally to the study.

## Abstract

Inflammatory arthritis and uveitis frequently co-exist, yet the mechanisms linking joint and ocular inflammation remain ill-defined. Here, we investigated how inflammatory arthritis influences ocular immune homeostasis using murine models of antigen-induced arthritis and collagen-induced arthritis. Arthritis promoted the accumulation of T cells, myeloid cells, and neutrophils within the vitreoretinal compartment, without progression to overt clinical uveitis. Ocular leukocyte recruitment was dynamically coupled to arthritis activity, resolving with remission of joint inflammation and recurring during arthritic flares. The magnitude of ocular immune perturbation correlated with arthritis severity, being enhanced in IL-27R-deficient mice and markedly reduced in IL-6R-deficient mice. Mechanistically, arthritis increased blood-retinal barrier permeability, demonstrating that systemic inflammation perturbs ocular immune privilege even in the absence of apparent ocular disease. While arthritis alone was insufficient to induce uveitis, it established a permissive ocular microenvironment that selectively enhanced the recruitment of adoptively transferred uveitogenic CD4^+^ T cells. These findings identify inflammatory arthritis as a systemic driver of subclinical ocular immune dysregulation and reveal a mechanism by which inflammation at a distant site may promote vulnerability to ocular autoimmunity. These data provide a framework for understanding immune dysregulation at the joint-eye axis and highlight cytokine pathways that may be targeted to preserve ocular immune homeostasis.

## 1. Introduction

The incidence of autoimmune and immune-mediated inflammatory diseases (IMIDs) is increasing, and their co-existence as multimorbidities presents an emerging global health challenge [1, 2]. A strong clinical association exists between sight-threatening uveitis and inflammatory arthritis [3, 4]. Uveitis and arthritis co-exist in approximately 30% of patients with spondyloarthropathies such as ankylosing spondylitis (AS) and psoriatic arthritis (PsA), with the highest prevalence observed in HLA-B27-positive disease [5]. Also at risk of concurrent joint and ocular inflammation are individuals with Behçet’s syndrome, children with PsA, and approximately 11-30% of children with juvenile idiopathic arthritis (JIA), who undergo regular screening to aid the early diagnosis and treatment of ocular involvement [6, 7]. The development of arthritis-associated uveitis in some patients highlights the heterogeneous nature of these conditions and a vulnerability to pathogenic mechanisms involving an interplay between inflamed joints and eyes.

While inflammatory arthritis and uveitis are typically studied separately, there is increasing evidence of shared immunological mechanisms that promote joint and ocular pathology. Cytokine-targeting drugs have emerged as important treatments for inflammatory arthritis and autoimmune uveitis, underscoring pathogenic roles for key cytokines. Tumor necrosis factor (TNF) blockers (e.g., adalimumab, infliximab) are frontline drugs for rheumatoid arthritis (RA), PsA, AS, and JIA, and are also approved for autoimmune uveitis [8–10]. The interleukin-(IL)-6 receptor (IL-6R) is similarly targeted (e.g., tocilizumab) in the clinic for treating inflammatory arthritis and autoimmune uveitis [11–13]. More recently, an improved understanding of pathogenic CD4^+^ T cells that secrete IL-17 has led to the approval of drugs targeting the T helper (Th)17 axis (e.g., IL-17, IL-23) in PsA and AS [14], with pre-clinical and early clinical investigations also revealing pathogenic roles for these cytokines in autoimmune uveitis [15–20]. Evidence supporting shared immunological mechanisms underlying arthritis and uveitis as comorbid conditions is illustrated by the observation that anti-TNF and other cytokine-targeting therapies, while significantly improving rheumatic symptoms in diseases such as JIA and spondyloarthropathies, have also been associated with reduced incidence of uveitis in these patients [6, 7, 21]. Recent clinical trials further highlight the potential of targeting the cytokines TNF and IL-6 for the management of arthritis-associated uveitis [22–24]. These findings suggest that common pathogenic pathways may drive inflammation in both joints and eyes. A deeper understanding of the immunological mechanisms shared between these tissues has the potential to identify early biomarkers for ocular involvement and novel therapeutic targets for treating uveitis in the context of inflammatory arthritis.

Animal models provide an opportunity to understand early ocular immune changes coupled to arthritis, which are challenging to capture in human disease. Indeed, pre-clinical models have illustrated a link between joint inflammation and ocular immune homeostasis [25–28]. In this study, we have utilized established models of inflammatory arthritis to gain insight into the relationship between joint inflammation and ocular immune homeostasis. We report that induction of arthritis is temporally coupled to disruption in blood-retinal barrier integrity and the recruitment of leukocytes to the eye. Using IL-27R- and IL-6R-deficient mice as models of severe and low disease activity respectively [29–33], we find that the degree of ocular leukocyte recruitment correlates with arthritis severity. These data provide new mechanistic insight into the relationship between joint inflammation and ocular homeostasis, and a route to exploring ocular tissue vulnerability during arthritis and clinical interventions that may maintain ocular homeostasis.

## 2. Materials and Methods

### 2.1. Mice

Wild-type (WT) C57BL/6 mice were purchased from Charles River UK. C57BL/6 mice deficient in the IL-6R (*Il6ra^-/-^*) lack exons 4-6 encoding the IL-6 binding motif and have been described previously [34]. C57BL/6 mice deficient in the IL-27R (B6N.129P2-*Il27ra^tm1Mak^*/J) were sourced from The Jackson Laboratory and have a targeted deletion of the *Il27ra* exon encoding the extracellular fibronectin type III domain, abolishing gene function [35]. The congenic C57BL/6 Ly5.1 (CD45.1^+^) strain carries the SJL mouse allele for the *Ptprc (Cd45)* gene and was originally obtained from The Jackson Laboratory. Heterozygous *Cx3cr1^CreER/+^*:*Rosa26^tdTomato^*^/+^ mice on a C57BL/6J background were used to label retinal microglia and were originally provided by Clemens Lange (University of Freiburg, Germany). To activate local and selective tdTomato expression in retinal microglia, eye drops were administered 3 times daily (every 2–3 h) with 5 mg/mL tamoxifen (T5648; Sigma-Aldrich, Poole, UK) in corn oil (C8267; Sigma-Aldrich) for 3 days, followed by a washout period of at least 4 weeks before experiments [36]. Male DBA/1 mice were purchased from Envigo UK. Mice were bred and housed at the University of Bristol Animal Services Unit and Cardiff University under specific pathogen-free conditions with food and water *ad libitum*. All procedures were approved by Ethical Review Groups at the University of Bristol and Cardiff University. Work was conducted in accordance with the UK Home Office PPL licenses PB3E4EE13, PF34A3DC8, PP0907754, PE8BCF782, PP3787461 and PP8830147.

### 2.2. Antigen-induced arthritis

Antigen-induced arthritis (AIA) was established in 8-10 week-old mice as described previously [29, 30, 33, 37]. Mice were immunized (s.c.) with methylated bovine serum albumin (mBSA; 1 mg/ml emulsified in Complete Freund’s Adjuvant) and 160 ng *Bordetella pertussis* toxin (i.p.). One week later, mice received an identical challenge of mBSA/CFA (s.c.). Inflammatory arthritis was established by intra-articular (i.a.) administration of mBSA (10μl; 10 mg/ml) into the right knee joint 21 days after the first mBSA/CFA immunization. Arthritis development was followed by measuring knee joint diameters using a POCO2T micrometer (Kroeplin) and eyes were harvested at 3 and 10 days following arthritis induction for assessment of ocular leukocyte infiltrates. Some mice received up to three flares of arthritis as described in the text by repeat administration of mBSA into the right knee joint (i.a.) [30].

### 2.3. Collagen-induced arthritis

Collagen-induced arthritis (CIA) was established as described previously [30, 37]. Briefly, 8-10 week-old DBA-1 male mice were immunized by intradermal injection (i.d.) with chick collagen type II (CII, 2 mg/ml; Sigma-Aldrich) dissolved in 10 mM acetic acid and emulsified with Incomplete Freund’s Adjuvant (Sigma-Aldrich) supplemented with 5 mg/ml Mycobacterium tuberculosis H37Ra (BD Biosciences). Each animal received an identical intradermal injection 21 days later. Arthritis severity was scored in each paw using a scale ranging from 0 to 4, as follows: 0, no evidence of erythema and swelling; 1, erythema and mild swelling confined to the tarsals or ankle joint; 2, erythema and mild swelling extending from the ankle to the tarsals; 3, erythema and moderate swelling extending from the ankle to metatarsal joints; 4, erythema and marked swelling encompass the ankle, foot, and digits. The aggregate values for each animal gave the clinical score.

### 2.4. Experimental autoimmune uveitis

Active experimental autoimmune uveitis (EAU) was induced as described previously using the Interphotoreceptor Retinoid-Binding Protein (IRBP) antigen, retinal binding protein RBP-3_629-643_, which was prepared by dissolving the peptide in 2% (v/v) DMSO in PBS at 20 mg/ml [38]. Female C57BL/6 mice at 8-10 weeks of age were immunized (s.c.) with 100 µl of 400 µg of RBP-3_629-643_ in CFA supplemented with 1.5 mg/ml *Mycobacterium tuberculosis.* At the same time, mice received a 100 µl injection (i.p.) of 1.5 µg/ml *Bordetella pertussis* toxin.

EAU was also established through the adoptive transfer of uveitogenic immune cells [39]. On day 11 following induction of EAU (before clinical disease onset), mice were killed and spleen and lymph nodes were recovered. Single cell suspensions were prepared through mechanical dissociation and passing through a 40 µm cell strainer into Dulbecco’s Modified Eagle Medium (DMEM). Cells were cultured (75 x 10^6^ cells per T75 flask) in DMEM supplemented with 10% (v/v) FBS, 2 mM glutamine, 100 U/ml penicillin, 100 µg/ml streptomycin, 10 µg/ml RBP-3_629-643_ and 10 ng/ml IL-23. Cells were cultured at 37°C, 4% CO2 for 72 hours, with 10 ng/ml IL-2 added after 24 hours. Cells were collected and washed with DMEM before isolation of mononuclear cells by density gradient centrifugation with Ficoll at 400 xg for 20 minutes. Cells were resuspended in PBS and the stated number of mononuclear cells injected (i.p.) into recipient mice. Changes in the ocular immune microenvironment were monitored by OCT and flow cytometry.

### 2.5. Ocular imaging

Pupils were first dilated with 1% (w/v) tropicamide and 2.5% (w/v) phenylephrine drops (Minims, Chauvin Pharmaceuticals), and mice anaesthetized with 2% isoflurane (Covetrus), with Viscotears (Novartis Pharmaceuticals) applied to protect the corneal surface. Retinal fundal images and OCT scans were captured on the Micron IV retinal imaging microscope (Phoenix Research Laboratories. Pleasanton, CA). Gain was set to +4 and the frames per second (FPS) to 15. Optical coherence tomography (OCT) images were standardized by taking one circle scan centered around the optic disc, with two full-length vertical and horizontal scans passing through the optic disc. Final images were averaged from 30 scans taken in rapid succession to enhance image quality. For tdTomato imaging, 550/25 nm band-pass excitation and 590 nm long-pass emission filters were applied (Edmund Optics, Barrington, NJ). Fundal and OCT images were assessed using the grading system described by Shome *et al*. [40]. However, ocular disturbances during AIA resulted in immunological changes that did not lead to clinical-grade uveitis.

### 2.6. Assessment of microvascular permeability

The Evans blue azo dye binds strongly to serum albumin, allowing visualization of blood-retinal barrier breakdown. To evaluate retinal microvascular permeability, 100 μl of 2% (w/v) Evans Blue dye (Sigma-Aldrich) was administered to mice by intravenous (i.v.) injection. Mice were killed 10 minutes later by a rising concentration of CO_2_ gas. Eyes were removed and immersed in fresh 2% (w/v) paraformaldehyde for 2 hours. Retinal whole mounts were prepared by removing the anterior segment and peeling the retina away from the choroid. Retinas were washed twice in cold PBS for 15 minutes, spread on clean glass slides and mounted, vitreous side up, under coverslips with mounting medium (Fluoroshield; Abcam). Whole mounts were examined for Evans blue (emission peak 680 nm) under a laser scanning confocal imaging system fitted with a multi-line argon laser (TCS-SP5-II-AOBS; Leica Microsystems). Quantification of Evans blue intensity and area was calculated using the “EBImage” and “RBioFormats” R packages. A mask of Evans blue signal was created and the average fluorescent intensity within this mask calculated. A mask containing the entire retinal tissue was created and the area of Evans blue staining was determined as a percentage of the total retinal tissue.

### 2.7. Histology

Knee joints were fixed in 10% (v/v) neutral buffered formalin and decalcified in 10% (v/v) formic acid at 4°C before embedding in paraffin. Parasagittal serial sections of 7 μm were stained with H&E, Safranin O and Fast Green (Sigma-Aldrich). Two independent observers, blinded to the experimental groups, scored the sections for subsynovial inflammation (0 = normal to 5 = ablation of adipose tissue due to leukocyte infiltrate), synovial exudate (0 = normal to 3 = substantial number of cells with large fibrin deposits), synovial hyperplasia (0 = normal 1–3 cells thick to 3 = over 3 layers thick with overgrowth onto joint surfaces with evidence of cartilage/ bone erosion), and cartilage/bone erosion (0 = normal to 3 = destruction of a significant part of the bone). The aggregate score is presented as an arthritic index.

### 2.8. Flow cytometry

Eyes were enucleated and dissected to recover the retina and vitreous in a 150 µl set volume of ice-cold PBS. The tissue was mechanically dissociated and passed through a 40 µm mesh 96-well filter plate to generate single-cell suspensions. Supernatant was discarded before resuspension of cells in FACS buzer (0.5% (w/v) bovine serum albumin, 5 mM ethylenediaminetetraacetic acid in D-PBS) containing rat anti-mouse CD16/CD32 Fc block (clone 2.4G2, BD Biosciences). Cells were then stained with fluorochrome-conjugated antibodies (Supplemental Table 1) specific to cell surface markers for 30 minutes at 4°C. Cells were acquired fresh without fixation following the addition of 7AAD for discrimination of live cells. To calculate absolute cell counts, a standard curve of seven 2-fold serial dilutions of a known concentration of splenocytes (top standard of 20 x 10^6^) was acquired on the flow cytometer in the same manner as the samples [41]. BD LSRII analyzer and BD Fortessa X20 flow cytometers were used to acquire samples, and analysis was completed using FlowJo software.

### 2.9. Statistics

Statistical comparisons of discontinuous histopathological scoring data involved the Mann Whitney U-test for comparison of two groups, or the Kruskal-Wallis test with Dunn’s multiple comparisons test for analysis of more than two groups. For analysis of flow cytometry data, a two-tailed unpaired *t-*test was used for comparison of two groups. For analysis of more than two groups, where significant differences were observed by one-way ANOVA, the Holm-Šidák test was used for multiple comparisons between groups. For analysis of CIA clinical data, a mixed-effects model with Šidák multiple correction was used. Data in graphs are expressed as mean ±SD. Analysis was conducted using Prism 10 (GraphPad Software). Values showing \**P*< 0.05; \*\**P*< 0.01; \*\*\**P*< 0.001, \*\*\*\**P*< 0.0001 were considered significant.

## 3. Results

### 3.1. Induction of inflammatory arthritis disrupts ocular immune homeostasis

To investigate how inflammatory arthritis influences the ocular immune microenvironment, antigen-induced arthritis (AIA) was established in wild-type (WT) male C57BL/6 mice. An increase in right knee (stifle) joint diameters relative to the contralateral control joint was observed in all mice (**Figure 1A**) and histopathology confirmed the classical hallmarks of inflammatory arthritis, including synovial leukocyte infiltration, synovial exudate, synovial hyperplasia, and bone and cartilage erosion (**Figure 1B**). Two time points were selected to examine arthritis-associated ocular immune alterations: the early peak of joint inflammation (2-3 days after intra-articular mBSA challenge) and a later, chronic-like phase (day 9-10) enriched in synovial T cell infiltrates and when joint swelling is resolving [30]. Fundal imaging showed no overt signs of ocular pathology (**Figure 1C**). However, OCT imaging at day 9 suggested a low degree of vitreoretinal cellular infiltrates in a subset of eyes (4 out of 8 mice), suggestive of a subclinical disruption in ocular homeostasis (**Figure 1C**). Indeed, flow cytometry of the vitreous and retina provided greater sensitivity, revealing significantly increased ocular leukocyte numbers at day 3 and day 10 of AIA compared to naïve control mice (**Figure 1D-E**), including elevated numbers of CD4^+^ and CD8^+^ T cells, CD11b^+^ myeloid cells and CD11b^+^Ly6G^+^ neutrophils (**Figure 1F**).

**Figure 1.**
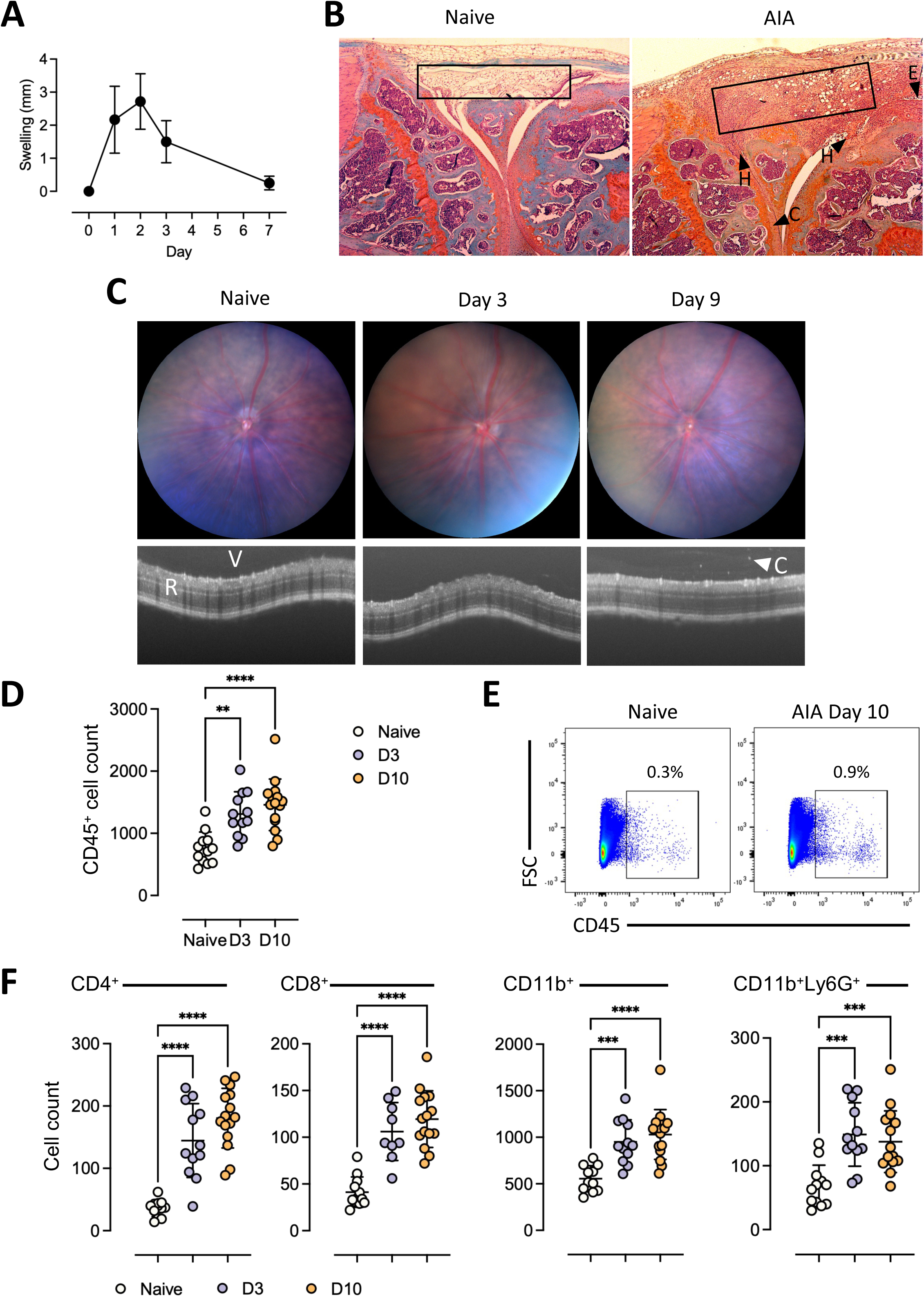
AIA leads to subclinical alterations in the ocular immune microenvironment. **(A)** Knee joint swelling calculated as the increase in joint diameters of the right arthritic knee relative to the contralateral control knee. **(B)** Representative histopathology at 5x magnification showing synovial infiltration (boxed area), synovial exudate (E arrowhead) and hyperplasia (H arrowhead), and cartilage erosion (C arrowhead) in mice with AIA compared to naïve control mice at day 10. **(C)** Representative fundal and OCT imaging at day 3 and 9 of AIA (R, retina; V, vitreous; C arrowhead, cellular infiltrate). **(D-E)** Number (D) and representative flow cytometry plots (E) of the CD45^+^ vitreoretinal cellular infiltrate at day 3 and day 10 of AIA compared to naïve eyes (n = 12-15 eyes/group). **(F)** Flow cytometric quantification of ocular CD4^+^ T cells, CD8^+^ T cells, CD11b^+^ cells and CD11b^+^Ly6G^+^ neutrophils at day 3 and day 10 of AIA compared to naïve eyes (n = 9-15 eyes/group). Graphs show mean ±SD analyzed by one-way ANOVA with Holm-Šídák multiple comparisons. **P<0.01, ***P<0.001, ****P<0.0001.

To determine whether these ocular immune changes were unique to the AIA model, we also established collagen-induced arthritis (CIA) in male DBA-1 mice. CIA onset was observed in 100% of mice, beginning at day 24 following the first CII challenge (**Figure 2A**). Disease severity peaked at day 30 with a mean clinical score of 8, individual paw score of 2, and an increase in paw swelling from approximately 2.2 to 2.6 mm (**Figure 2B-D**). Consistent with that observed in AIA, CIA was accompanied by a significant accumulation of ocular leukocytes, comprising CD4^+^ T cells, CD8^+^ T cells and CD11b^+^Ly6G^+^ neutrophils (**Figure 2E**). Together, these findings demonstrate that both AIA and CIA disrupt ocular immune homeostasis, marked by leukocyte accumulation in the eyes.

**Figure 2.**
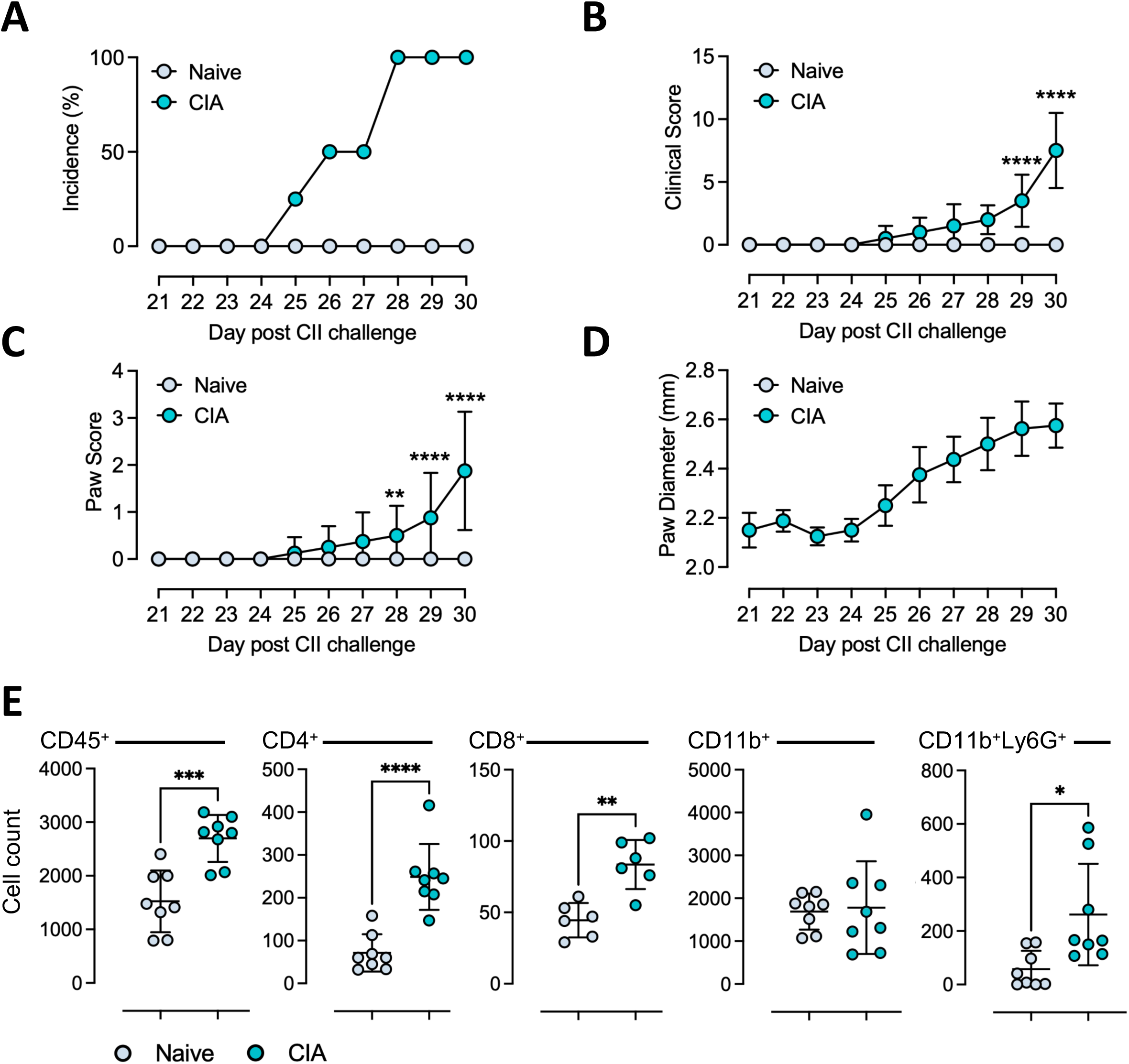
CIA promotes an increase in ocular leukocyte recruitment. **(A-D)** Incidence (A), clinical score (B), average paw scores (C) and hind paw diameters (D) of mice with CIA from day 21-30 compared to naïve control mice (n = 4 mice/group). **(E)** Flow cytometric quantification of ocular CD45^+^ cells, CD4^+^ T cells, CD8^+^ T cells, CD11b^+^ cells and CD11b^+^Ly6G^+^ neutrophils at day 30 of CIA compared to naive mice (n = 6-8 eyes/group). Graphs show mean ±SD analyzed by mixed-effects model with Šidák multiple correction (B, C) and unpaired t test (E). *P<0.05, **P<0.01, ***P<0.001, ****P<0.0001.

### 3.2. Ocular leukocyte recruitment coincides with flares of joint inflammation

A temporal association between joint and ocular inflammation has been reported in arthritis-associated uveitis, suggesting that clinical flares of arthritis and uveitis occur concurrently [42]. We therefore questioned whether the increase in ocular leukocytes in AIA followed the same temporal pattern as joint inflammation, and whether recurrent flares of arthritis would lead to cumulative ocular leukocyte infiltration resulting in an exacerbation and clinical signs of uveitis. To this end, AIA was established in WT male C57BL/6 mice, and three flares of joint inflammation were triggered by repeat intra-articular administration of mBSA at day 0 (flare 1), day 14 (flare 2) and day 28 (flare 3), as previously described for modelling relapsing–remitting arthritis [30] (**Figure 3A**). Measurement of inflamed knee joint diameters relative to the contralateral control joint confirmed that swelling followed a consistent temporal pattern across each flare, involving an early peak in joint inflammation (1-3 days post-flare induction) followed by a period of resolution (3-14 days post-flare induction) (**Figure 3B**). OCT and fundal signs imaging during each flare revealed that multiple episodes of arthritis did not lead to overt clinical signs of uveitis. Instead, a comparable level of subclinical leukocyte infiltration was observed across each flare (**Figure 3C**). Imaging showed no evidence of leukocyte infiltration in mice systemically primed to generate an mBSA-specific immune response in the absence of intra-articular mBSA administration (Primed - no arthritis group). Leukocyte infiltrates were also undetectable by OCT imaging once joint swelling had subsided (Resolved group).

**Figure 3.**
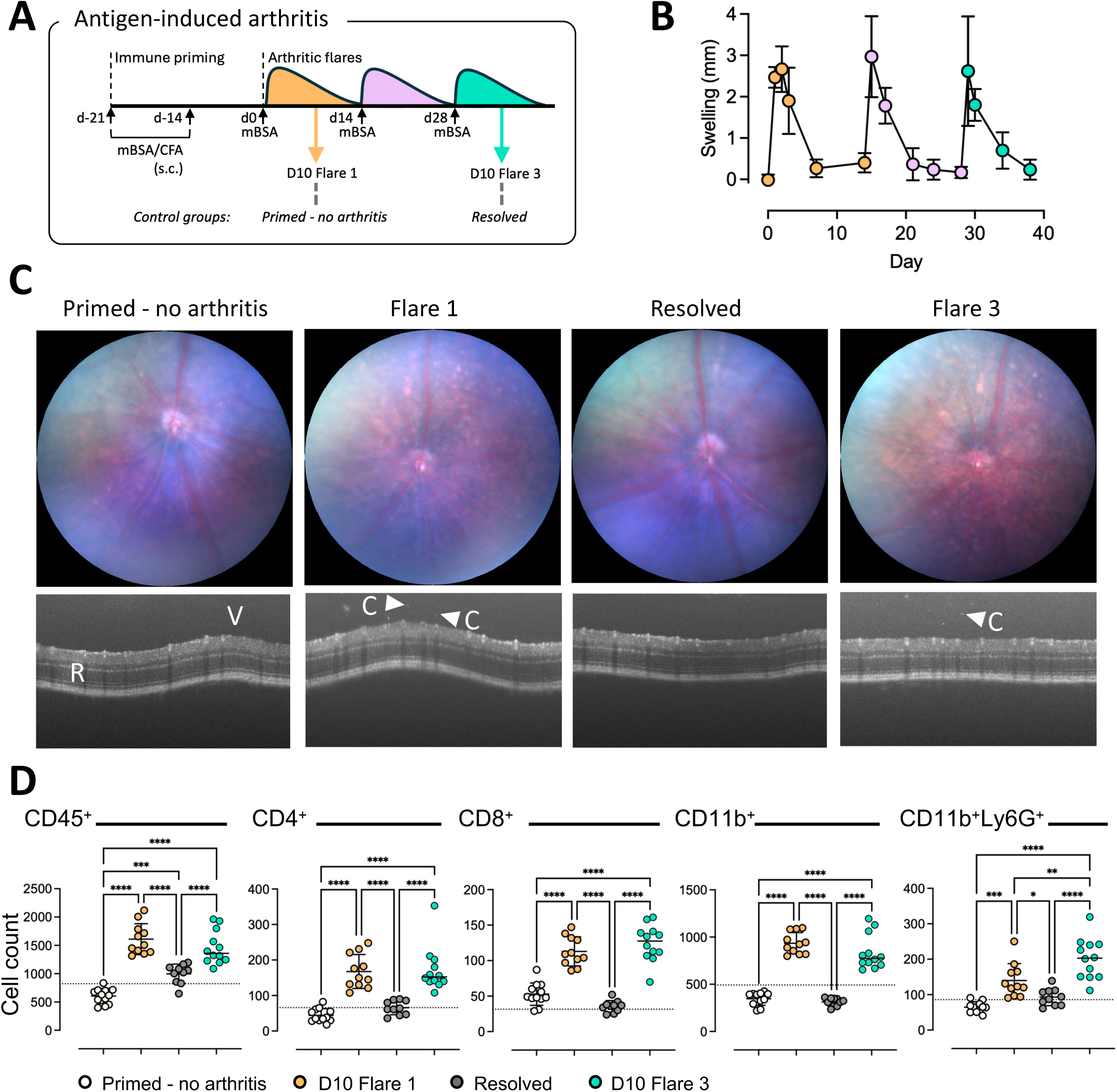
Ocular leukocyte recruitment temporally mirrors the pattern of arthritic flares in AIA. **(A)** Schematic demonstrating the time points used to assess ocular recruitment in a recurrent model of AIA. Ocular leukocyte numbers were quantified following systemic priming for AIA with mBSA/CFA (Primed - no arthritis), at day 10 of the first arthritic flare (D10 Flare 1), 24 days after induction of two flares of arthritis and resolution of joint swelling (Resolved) and at day 10 during the third flare of arthritis (D10 Flare 3). **(B)** Knee joint swelling, measured as the increase in joint diameters of the right arthritic knee relative to the contralateral control joint was used to track the three recurrent episodes of joint inflammation. **(C)** Representative fundal and OCT imaging in primed mice, at day 10 of Flare 1, following resolution of joint swelling, and at day 10 of Flare 3 (R, retina; V, vitreous; C arrowhead, cellular infiltrate). **(D)** Flow cytometric quantification of ocular CD45^+^ cells, CD4^+^ T cells, CD8^+^ T cells, CD11b^+^ cells and CD11b^+^Ly6G^+^ neutrophils in primed mice without arthritis induction, at day 10 Flare 1, following resolution of joint swelling, and at day 10 Flare 3 (n = 10-12 eyes/group). Dotted line represents mean cell number in control Naïve mice. Graphs show mean ±SD analyzed by one-way ANOVA with Holm-Šídák multiple comparisons. *P<0.05, **P<0.01, ***P<0.001, ****P<0.0001. mBSA, methylated bovine serum albumin; CFA, Complete Freund’s Adjuvant.

Consistent with that observed by OCT imaging, flow cytometry revealed a comparable degree of CD45^+^ cell, CD4^+^ and CD8^+^ T cell, CD11b^+^ myeloid cell and CD11b^+^Ly6G^+^ neutrophil cells in eyes at day 10 following induction of Flare 1 and Flare 3 of AIA (**Figure 3D**). Here, using *Cx3cr1^CreER/+^*:*Rosa26^tdTomato^*^/+^ mice to label microglia revealed a local expansion of these cells in the eye, contributing to the observed increase in CD11b^+^ myeloid cells (**Supplemental Figure 1**). Negligible changes in ocular leukocyte numbers were observed in mice that were only primed for an mBSA-specific immune response, confirming that arthritis induction was required to disrupt ocular immune homeostasis. Consistent with this observation, intra-articular administration of PBS also had negligible impact on ocular leukocyte numbers (**Supplemental Figure 2**). To determine whether the ocular leukocyte response resolved following arthritic flares, ocular leukocyte numbers were compared between mice at day 10 of Flare 3 and mice at the equivalent time point, but in which inflammation had resolved after receiving only two arthritic flares (Resolved group). A significant decrease in CD4^+^ and CD8^+^ T cells, CD11b^+^ myeloid cells and CD11b^+^Ly6G^+^ neutrophils was observed in the resolved group, indicating that ocular leukocyte accumulation resolves in parallel with joint inflammation. Collectively, these data demonstrate that increased ocular leukocyte numbers are dynamically coupled to episodes of joint inflammation in AIA. Despite repeated arthritic flares, disruption of ocular immune homeostasis did not accumulate over time to produce overt clinical uveitis.

### 3.3. The severity of arthritis determines the degree of ocular immune cell accumulation

Cytokine modulators are established therapies for inflammatory arthritis, autoimmune uveitis and arthritis-associated uveitis [8–13, 22–24]. To explore the relationship between disease activity and ocular leukocyte recruitment, we used mice deficient in IL-6 and IL-27 signaling to model opposing inflammatory states. In AIA, *Il27ra^-/-^* mice display severe arthritis [29], while IL-6 deficiency (*Il6^-/-^* and *Il6ra^-/-^* mice) results in protection [30, 31, 33, 43]. Similarly, in experimental autoimmune encephalitis (EAU), genetic deficiency or therapeutic blockade of IL-6 signaling limits ocular inflammation [44], whereas our analysis of *Il27ra^-/-^* mice reveals earlier onset and increased disease severity consistent with previous studies (**Supplemental Figure 3**) [18, 45]. Measurement of knee joint diameters during AIA confirmed an increase in swelling in *Il27ra^-/-^* mice compared to WT controls, while swelling in *Il6ra^-/-^*mice was significantly reduced and transient (**Figure 4A**). Histopathology confirmed that joint pathology in *Il6ra^-/-^* mice was significantly decreased compared to WT and *Il27ra^-/-^* mice (**Figure 4B-C**). Flow cytometry analysis of eyes at day 10 of AIA revealed an increase in CD45^+^ leukocytes in *Il27ra^-/-^* mice compared to WT controls, driven by elevated CD11b^+^ myeloid cells and CD11b^+^Ly6G^+^ neutrophils (**Figure 4D**). In contrast, *Il6ra^-/-^* mice showed significantly reduced numbers of CD45^+^ leukocytes, CD4^+^ and CD8^+^ T cells, CD11b^+^ myeloid cells and CD11b^+^Ly6G^+^ neutrophils compared to both WT and *Il27ra^-/-^* mice (**Figure 4D**). Here, ocular leukocyte numbers in *Il6ra^-/-^* mice were comparable with those in naïve WT controls. Together, these findings show that arthritis severity correlates with the degree of ocular immune cell accumulation, with IL-27 signaling acting as a negative regulator and IL-6 signaling promoting both joint inflammation and disruption of ocular immune homeostasis.

**Figure 4.**
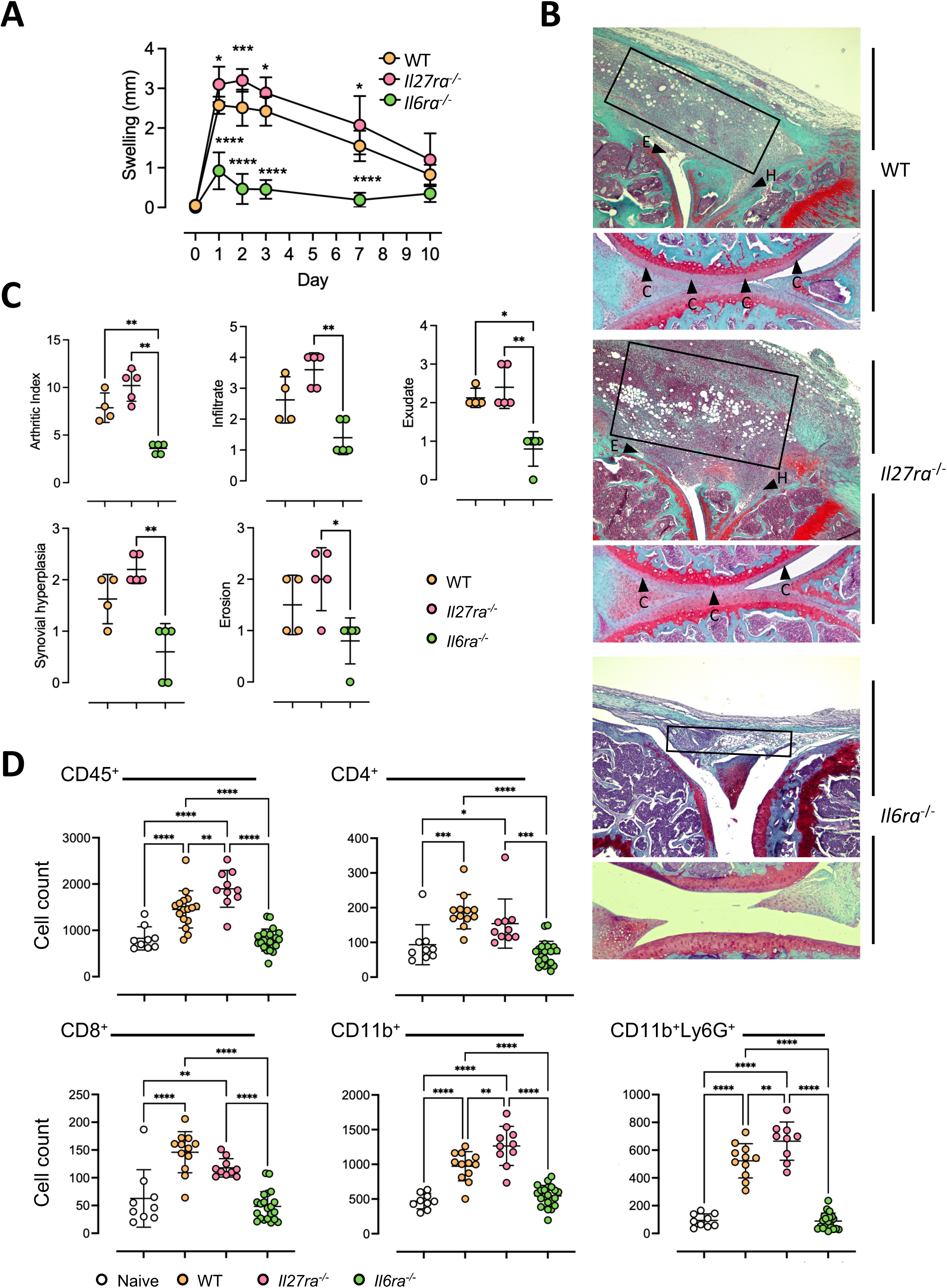
The magnitude of ocular leukocyte recruitment correlates with the severity of AIA. **(A)** Knee joint swelling, measured as the increase in joint diameters of the right arthritic knee relative to the contralateral control joint in male WT, *Il27ra^-/-^* and *Il6ra^-/-^* mice. Graph show mean ±SD analyzed by a mixed-effects model with Dunnett’s multiple correction to compare against the WT control group (n = 7-9 mice/group) **(B)** Representative histopathology at 5x magnification showing the extent of synovial infiltration (boxed area), synovial exudate (E arrowhead) and hyperplasia (H arrowhead), and cartilage erosion (C arrowhead) in WT, *Il27ra^-/-^* and *Il6ra^-/-^* mice at day 10 of AIA. **(C)** Histopathology scoring of WT, *Il27ra^-/-^* and *Il6ra^-/-^* mice at day 10 of AIA, showing arthritic index, synovial infiltration, exudate, hyperplasia and erosion (n = 4-5/group). Graphs show mean ±SD analyzed by Kruskal-Wallis with Dunn’s multiple correction. **(D)** Flow cytometric quantification of ocular CD45^+^ cells, CD4^+^ T cells, CD8^+^ T cells, CD11b^+^ cells and CD11b^+^Ly6G^+^ neutrophils in Naïve control mice and WT, *Il27ra^-/-^* and *Il6ra^-/-^*mice at day 10 of AIA (n = 9-21 eyes/group). Graphs show mean ±SD analyzed by one-way ANOVA with Holm-Šídák multiple correction. *P<0.05, **P<0.01, ***P<0.001, ****P<0.0001.

### 3.4. AIA compromises blood-retinal barrier integrity in the absence of overt uveitis

Our data reveal that induction of inflammatory arthritis alters the ocular immune microenvironment. Comparison of AIA and EAU by OCT imaging highlights the marked difference in the severity of ocular disruption caused by uveitis compared to arthritis. While subclinical levels of infiltrating ocular leukocytes are observed at peak AIA (day 10), mice at peak EAU (day 21) display severe (grade 3-4) pathology featuring large aggregates of hyperreflective debris and disruption of multiple retinal layers (**Figure 5A**) [40, 46]. Consistent with these observations, flow cytometry also confirmed that the magnitude of ocular CD45^+^ leukocytes and CD4^+^ T cell infiltrate is significantly lower in AIA than in EAU (**Figure 5B**).

**Figure 5.**
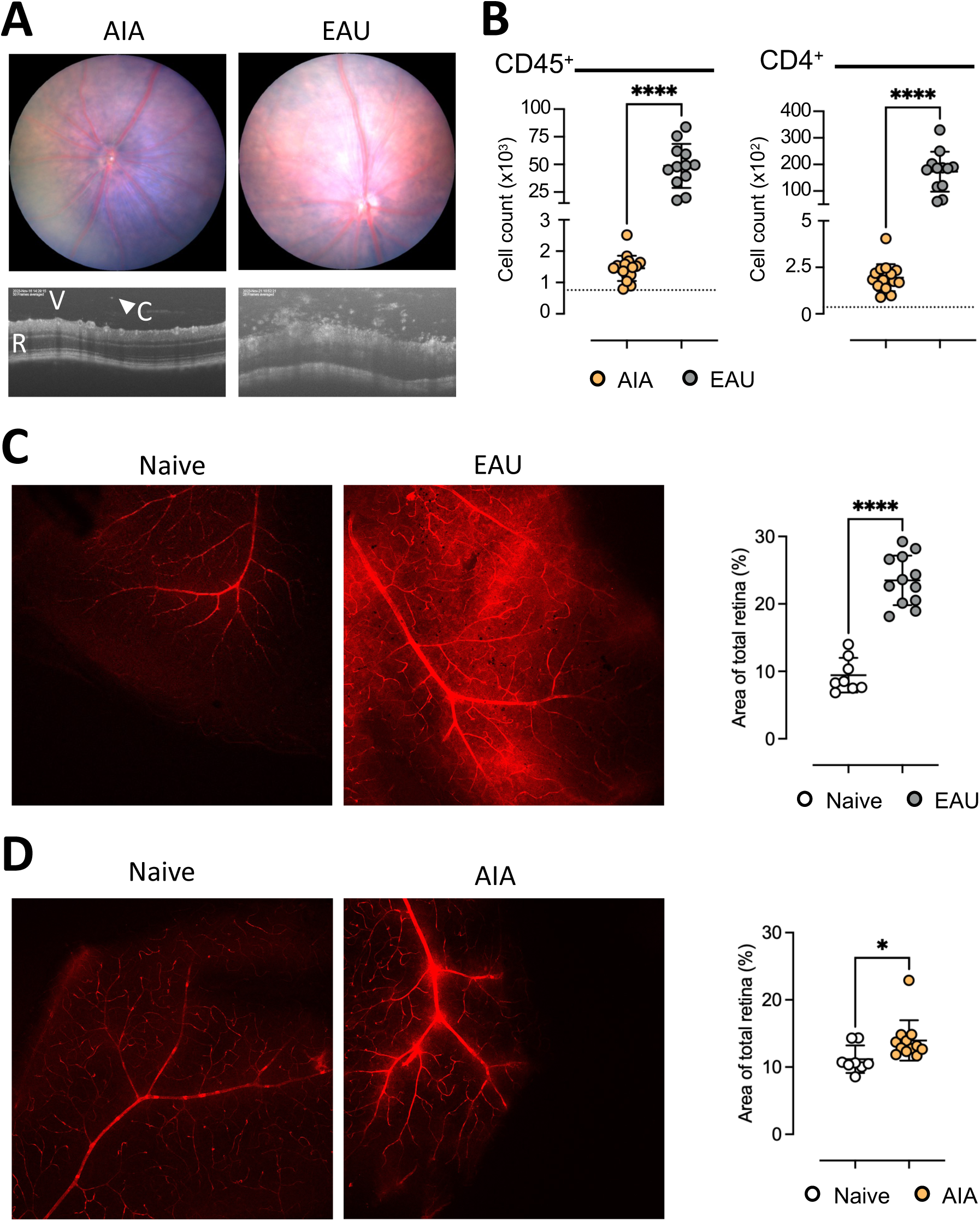
AIA leads to increased blood-retinal barrier permeability. **(A)** Representative fundal and OCT imaging of eyes at day 10 of AIA and day 21 of active EAU (R, retina; V, vitreous; C arrowhead, cellular infiltrate). **(B)** Quantification of CD45^+^ and CD4^+^ cells by flow cytometry comparing eyes from mice with AIA (n = 12 eyes) and EAU (n = 16 eyes). Dotted line represents mean cell number in control naïve mice. **(C-D)** Representative retinal flatmount images showing Evans Blue staining and quantification of staining intensity normalized by tissue area at day 21 of active EAU (n = 12 eyes) compared to naïve control eyes (n=8 eyes) (C) and at day 10 of AIA (n = 12 eyes) compared to naïve control (n = 8 eyes) (D). Graphs show mean ±SD analyzed by unpaired t test. *P<0.05, ****P<0.0001.

The blood-retinal barrier (BRB) regulates molecular transport and maintains ocular immune privilege to preserve visual function [47, 48]. Given that BRB disruption is a hallmark of uveitis and facilitates leukocyte infiltration, we next questioned whether subclinical ocular inflammation observed during AIA is associated with altered vascular permeability. As expected, assessment of microvascular permeability in retinal wholemounts during EAU revealed substantial Evans Blue leakage from the retinal vasculature, consistent with blood-ocular barrier breakdown (**Figure 5C**). Notably, mice with AIA also exhibited a significant increase in Evans Blue leakage compared to naïve control mice, although to a lesser extent than in EAU (**Figure 5D**), which is consistent with the lower degree of ocular leukocyte involvement observed during AIA. These findings suggest that even subclinical ocular inflammation observed in AIA is accompanied by increased microvascular permeability, suggesting that arthritis may contribute to an early compromise of BRB integrity.

### 3.5. AIA supports ocular recruitment of uveitogenic T helper cells

In inflammatory arthritides such as JIA, arthritis onset typically precedes the development of uveitis [49–51]. We, therefore, investigated whether the disruption of ocular immune homeostasis observed during AIA may increase vulnerability to uveitis. To address this, the AIA model was combined with a model of adoptive transfer of EAU, in which pathogenic leukocytes from mice with EAU were transferred into mice with AIA [37, 39]. Here, our previous work has shown that the transfer of 2 x 10^6^ leukocytes from EAU-primed mice induces disease in >80% of recipient mice, and that CD4^+^ T cells are both necessary and sufficient to induce uveitis [39].

Uveitogenic leukocytes were recovered from the spleen and lymph nodes of mice immunized for EAU and stimulated *ex vivo* with RBP-3_629-643_, IL-23 and IL-2, before transfer into recipient mice at day 10 of AIA (**Figure 6A**). Consistent with our previous findings, the transfer of 1 x 10^6^ leukocytes into recipient male mice resulted in robust clinical uveitis, and a comparable degree of ocular leukocyte recruitment in both mice with and without AIA (**Supplemental Figure 4**). Given the substantial disease observed following transfer of 1 x 10^6^ leukocytes, we next transferred a ‘subthreshold’ number of leukocytes (2 x 10^5^), below that required to establish EAU, to determine whether AIA enhanced the ocular accumulation of leukocytes with pathogenic potential that may increase vulnerability to clinical uveitis. Flow cytometry revealed comparable numbers of total CD45^+^ cells, CD8^+^ T cells, and CD11b^+^ myeloid cells in the eyes of mice with and without AIA (**Figure 6B**). However, mice challenged with AIA showed increased numbers of ocular CD4^+^ T cells, which was predominantly driven by enhanced recruitment of the transferred pathogenic CD45.1^+^ cells, rather than endogenous CD45.2^+^ CD4^+^ T cells. Although AIA studies are traditionally performed in male mice, female mice also display high susceptibility to AIA. We therefore assessed whether the transfer of uveitogenic leukocytes into female mice similarly increased ocular CD4^+^ T cell accumulation. Here, while the total number of ocular CD4^+^ T cells was not significantly increased in female mice with AIA, the number of transferred pathogenic CD45.1^+^ CD4^+^ T cells was significantly elevated compared to control mice (**Figure 6C**). These findings indicate that inflammatory arthritis potentiates the ocular recruitment of pathogenic CD4^+^ T cells.

**Figure 6.**
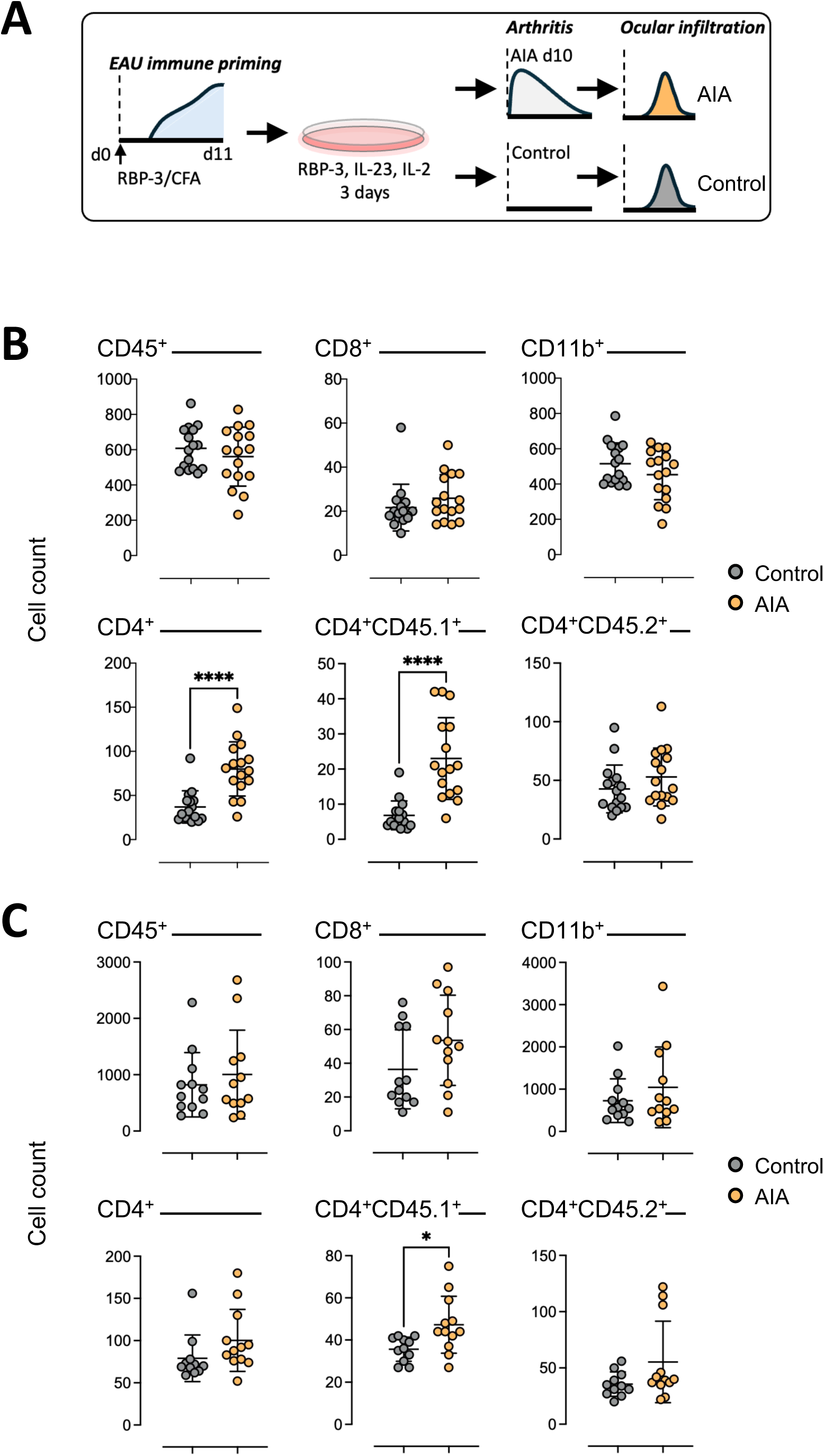
AIA supports the selective recruitment of uveitogenic CD4^+^ T cells. **(A)** Schematic of the experimental approach involving the adoptive transfer of uveitogenic cells from EAU-primed donor female C57BL/6 Ly5 (CD45.1^+^) mice into C57BL/6 WT (CD45.2^+^) recipient mice. Prior to transfer, leukocytes were cultured for 3 days in the presence of RBP-3_629-643_ and IL-23, with IL-2 added after the first 24 h of culture. Leukocytes were transferred into control mice without AIA, or mice at day 10 of AIA. **(B)** Flow cytometric quantification of ocular CD45^+^ cells, CD8^+^ T cells and CD11b^+^ myeloid cells and the total CD4^+^ T cells, transferred CD45.1^+^ CD4^+^ T cells and endogenous CD45.2^+^ CD4 T cells. Graphs compare male mice with AIA and control mice (n = 16 eyes per group) at day 11 post transfer of uveitogenic leukocytes. **(C)** Flow cytometric quantification of ocular CD45^+^ cells, CD8^+^ T cells and CD11b^+^ myeloid cells and the total CD4^+^ T cells, transferred CD45.1^+^ CD4^+^ T cells and endogenous CD45.2^+^ CD4^+^ T cells. Graphs compare female mice with AIA and control mice (n = 11-12 eyes per group) at day 11 post transfer of uveitogenic leukocytes. Graphs show mean ±SD analyzed by unpaired t test. *P<0.05, ****P<0.0001

## 4. Discussion

A strong clinical association exists between inflammatory arthritis and uveitis, yet understanding of the immunological relationship linking co-existing joint and ocular pathology remains incomplete. In this study, we offer new insights into how inflammatory arthritis impacts the ocular immune microenvironment. Induction of arthritis was sufficient to perturb ocular immune homeostasis, leading to transient leukocyte accumulation in the vitreoretinal compartment and increased BRB permeability in the absence of overt clinical uveitis. Combining complementary models of antigen-induced arthritis, collagen-induced arthritis and experimental autoimmune uveitis, we demonstrate that this phenomenon is robust, temporally coupled to episodes of joint inflammation, and quantitatively linked to the severity of arthritis. Collectively, our data identify inflammatory arthritis as a systemic driver of ocular immune perturbation, with important implications for local immunosurveillance at an immune-privileged site.

Uveitis is a term for over 30 conditions involving intraocular inflammation, a significant proportion of which are associated with IMIDs. Rheumatological conditions feature prominently among comorbidities associated with uveitis, and include the spondyloarthropathies, Behçet’s syndrome, Kawasaki disease, sarcoidosis, Sjögren’s syndrome, reactive arthritis, and JIA [3, 4]. Arthritis frequently precedes uveitis onset in JIA, suggesting that systemic immune activation arising outside the eye creates conditions permissive for ocular immune dysregulation [49–51]. Our data support this concept by demonstrating that inflammatory arthritis alone, without concurrent induction of ocular autoimmunity, perturbs ocular immune privilege by facilitating leukocyte recruitment to the eye.

Previous experimental models have highlighted links between arthritis and ocular inflammation. The SKG mouse model, driven by a mutation in the zeta-chain-associated protein kinase 70 (*Zap70*) gene, results in defective thymic selection and the escape of autoreactive T cells that promote spontaneous arthritis [52]. In this model, synchronization of arthritis onset through the administration of fungal glucans drives a polyarthritis involving extraarticular manifestations reminiscent of the spondyloarthropathies, including uveitis [28]. Similarly, studies utilizing intravital videomicroscopy in the SKG model describe ocular intravascular cell trafficking and leukocyte-endothelial interactions coinciding with arthritis onset, which does not lead to the development of overt uveitis [25]. In a T cell receptor-transgenic (TCR-Tg) mouse model of arthritis, recognizing a dominant arthritogenic epitope of cartilage proteoglycan, mice also displayed increased cellular rolling and adhesion in the iris vasculature, and increased cellular infiltration within the eye associated with arthritis [27]. These observations suggest an increase in microvascular permeability and endothelial activation within the eye following arthritis induction. Our data support and extend these observations by demonstrating increased ocular vascular permeability during AIA and by defining the immune composition of the arthritis-associated ocular leukocyte population. Although the degree of BRB disruption observed during AIA was substantially less severe than in EAU, the increase in Evans Blue leakage demonstrates that joint inflammation alone is sufficient to compromise retinal vascular integrity. Consistent with this observation, OCT imaging revealed a low-level vitreoretinal infiltrate during arthritis, while flow cytometric analysis confirmed the accumulation of ocular leukocytes comprising CD4^+^ and CD8^+^ T cells, CD11b^+^ myeloid cells and CD11b^+^Ly6G^+^ neutrophils. Importantly, the magnitude of this response remained markedly lower than observed during overt autoimmune uveitis.

Induction of the monoarticular AIA model was sufficient to disrupt the ocular immune compartment and allowed us to model the temporal dynamics of arthritis-associated ocular leukocyte recruitment. In JIA, inflammatory markers including S100 proteins (S100A8/A9 and S100A12), IL-8, and sICAM in the serum and aqueous humour have been proposed as biomarkers of disease activity and uveitis risk [53–55]. Similarly in AS, patients with uveitis display increased antibodies specific to prefoldin-5, which have been proposed as biomarkers of ocular involvement [56]. The presence of prefoldin-5-specific antibodies were also observed in the SKG mouse model of inflammatory arthritis, where prefoldin-5 displayed a protective anti-apoptotic role within the eye [56]. Such studies combining the analysis of human cohorts with mechanistic studies in animal models have the potential to identify novel biomarkers of early disease activity and uveitis flares. A notable finding of our study is that ocular leukocyte recruitment mirrored the kinetics of joint inflammation, where recurrent flares of arthritis produced transient increases in ocular leukocytes that resolved alongside synovitis, and without cumulative progression to uveitis. These observations suggest that systemic inflammation may repeatedly challenge ocular immune privilege but is not, by itself, sufficient to trigger ocular disease. Instead, additional pathogenic signals may be required to establish disease, including local antigen recognition, activation of resident antigen-presenting cells, altered chemokine production and the failure of local regulatory mechanisms. This idea is supported by our adoptive transfer experiments, where AIA selectively enhanced the recruitment of pathogenic uveitogenic CD4^+^CD45.1^+^ T cells into the eye. Inflammatory arthritis may, therefore, establish a permissive ocular microenvironment that lowers the threshold for the infiltration or retention of uveitogenic lymphocytes.

Susceptibility to IMIDs including inflammatory arthritis is associated with CD4^+^ T cells, Th17 responses, IL-23 signaling and STAT3 activation [57, 58]. Targeting the Th17/IL-23 axis has emerged as an important treatment for the spondyloarthropathies [14], and is being explored in autoimmune uveitis [15–20]. AIA is also dependent on CD4^+^ T cells, Th17 cells and IL-6-mediated STAT3 signaling [30, 43, 59–61], and blockade of IL-6 signaling (e.g., tocilizumab) has advanced the treatment of inflammatory arthritis [11]. Consistent with these observations, a loss of IL-6 signaling in *Il6ra^-/-^* mice ameliorated AIA and mice showed a significant decrease in ocular leukocytes. Both IL-6 and IL-27 share common biology by signaling through the common β-signaling receptor, gp130, to activate STAT1 and STAT3. However, IL-6 and IL-27 often oppose each other’s actions, where IL-27 is a more potent STAT1 activator and can inhibit IL-6/STAT3-driven outcomes to support regulatory-like responses that limit CD4^+^ T cell-driven pathology [62–65]. While IL-6 promotes arthritis progression, IL-27 limits arthritis severity [29, 32]. Our studies revealed that *Il27ra^-/-^* mice displayed a significantly higher accumulation of ocular leukocytes during AIA compared to WT and *Il6ra^-/-^* mice, suggesting that the overall inflammatory burden of arthritis is an important determinant of ocular immune dysregulation. While these observations do not distinguish between the direct effects of IL-6 and IL-27 within the eye and indirect effects mediated through modulation of arthritis severity, they support a model in which systemic cytokine networks influence ocular immune homeostasis. The opposing roles of IL-6 and IL-27 in regulating STAT3- and STAT1-dependent immune responses may therefore represent an important axis controlling susceptibility to ocular immune perturbation during arthritis. In this regard, blockade of IL-6 signaling via its membrane-bound IL-6R (classical IL-6 signaling) or the soluble form of the IL-6R (IL-6 trans-signaling) may offer potential to both limit the inflammatory responses and also prevent BRB breakdown [66–68]. In this regard, phase III clinical trials of an anti-IL-6R blocker (vamikibart) have shown early positive signs in uveitic macular edema [69].

The selective accumulation of uveitogenic CD4^+^ T-cells (CD45.1^+^) with pathogenic potential compared to endogenous CD4 T cells (CD45.2^+^) during AIA has important implications for understanding arthritis-associated uveitis. Th17-associated cytokines, including IL-17, IL-23 and IL-6 play established pathogenic roles in both inflammatory arthritis and autoimmune uveitis [14, 18–20, 29, 30, 33]. Our transferred cells were generated under conditions that support pathogenic Th17-like responses and we have previously shown these CD4^+^ T cells produce IL-17 and IFNγ [39]. These observations suggest that arthritis-induced disruption of ocular immune homeostasis preferentially facilitates the recruitment and/or persistence of activated pathogenic CD4^+^ T cells rather than indiscriminately increasing all leukocyte populations. Consequently, therapeutic approaches targeting pathways central to CD4/Th17 biology may offer benefits across inflammatory arthritis, autoimmune uveitis and arthritis-associated uveitis.

In summary, our findings demonstrate that inflammatory arthritis perturbs ocular immune privilege by promoting leukocyte recruitment and compromising BRB integrity. These changes are dynamically linked to arthritis severity and enhance ocular recruitment of pathogenic CD4^+^ T cells. Our data identify systemic inflammatory arthritis as a driver of subclinical ocular immune dysregulation and provide mechanistic insight into why some patients with inflammatory arthritis may be vulnerable to developing uveitis. More broadly, these data support the concept that IMIDs can alter the homeostasis of anatomically distant immune-privileged tissues, with implications for understanding comorbidity across chronic inflammatory disorders.

## Acknowledgements

Research was supported by grants from Fight for Sight (References Dr Gareth Jones_PhDa_20200610_026, Dr Gareth Jones_PG2023_20231030_003), Versus Arthritis (References 22706, 20305, 20770) and UKRI-MRC (Reference MR/X00077X/1) and The Underwood Trust. DGH and CM received a PhD Studentship from the GW4 BioMed UKRI-MRC Doctoral Training Partnership. AM was funded by an industry-sponsored PhD from Galecto Biotech. AW was funded by Sight Research UK (formerly National Eye Research Centre UK). DAC and JL were supported by the National Institute for Health and Care Research Biomedical Research Center (BRC) based at Moorfields Eye Hospital NHS Foundation Trust and the UCL Institute of Ophthalmology (BRC4-03-RB414-405). The authors wish to thank Andrew Herman, Helen Rice, Celyn Dugdale, (Flow Cytometry Facility, University of Bristol) and the University of Bristol Animal Services Unit for their technical assistance. For the purpose of Open Access, the authors have applied a Creative Commons Attribution (CC BY) licence to any Author Accepted Manuscript version arising from this submission.

## Author Contributions (CREdiT)-

*Conceptualization*: GWJ, LBN

*Data curation:* SE, AW, CM, DGH, AM, SD, AY, GWJ

*Formal analysis:* SE, AW, CM, DGH, AM, SD, AY, GWJ

*Funding acquisition:* SAJ, ADD, LBN, GWJ

*Investigation:* SE, AW, CM, DGH, AM, SD, AY, JL, DAC, GWJ

*Methodology:* SE, AW, CM, ASW, JL, ADD, DAC, LBN, GWJ

*Resources:* SAJ, ADD, DAC, LBN, GWJ

*Supervision:* SAJ, ADD, DAC, LBN, GWJ

*Visualization*: SE, AW, CM, AM, GWJ

*Writing – initial draft*: SE, AW, LBN, GWJ

*Review and editing:* All authors

---

**Supplemental Figure 1. Resident ocular microglia numbers increase after AIA induction**

AIA was established in *Cx3cr1^CreER^*^/+^:*Rosa26^tdTomato^*^/+^ mice to label microglia. **(A)** Representative fundus imaging for fluorescent detection of tdTomato-labelled CX3CR1-positive microglia in mice during 3 flares of arthritis showed no detectable change in phenotype such as perivascular sheathing. **(B)** Enumeration of CD45^+^CD11b^+^tdTomato^+^ cells by flow cytometry at day 10 of the first arthritic flare and day 10 of the third arthritic flare (n=11-12 eyes/group). Graphs show mean ±SD analyzed by one-way ANOVA with Holm-Šídák multiple comparisons. **P<0.01, ***P<0.001.

**Supplemental Figure 2. Intra-articular administration of PBS does not promote ocular leukocyte recruitment**

C57BL/6 mice were administered intra-articular PBS and ocular CD45^+^ leukocytes were quantified by flow cytometry after 10 days. Graph shows mean ±SD analyzed by unpaired t test.

**Supplemental Figure 3.. IL-27 signaling limits the severity of EAU.**

**(A)** *Il27ra^+/+^* (CD45.2) and *Il27ra^-/-^* (CD45.2) mice were immunized with RBP-3_629-643_. Flow cytometry of the vitreoretinal infiltrate for CD4^+^ and CD8^+^ T cells and CD11b^+^ myeloid cells at day 11 post challenge shows early onset of uveitis in *Il27ra*^-/-^ mice (n=8-10 per group). **(B-C)** At day 11 post challenge with RBP-3_629-643_, draining lymph nodes were recovered, re-stimulated with RBP-3_629-643_ in the presence of IL-2 and IL-23 for 72 hours, before adoptive transfer into recipient WT (CD45.1) mice. **(B)** OCT imaging shows exacerbated uveitis following the transfer of *Il27ra*^-/-^ cells compared to WT control (day 0-143). **(C)** Flow cytometry at day 67 post transfer shows that *Il27ra*^-/-^ cells promote increased infiltration and retention of both endogenous (CD45.1) and donor (CD45.2) CD4^+^ T cells in the eyes (n=9 per group) and an increase in ocular-infiltrating CD11b^+^Ly6G^+^ neutrophils (n=6 per group) at day 67 post transfer. Graphs show mean ± SD analyzed by unpaired t test (A), area under the curve, AUC (B) and one-way ANOVA with Holm-Šídák multiple comparisons (C). *P<0.05, **P<0.01, ***P<0.001, ****P<0.0001.

**Supplemental Figure 4. Adoptive transfer of 1 x 10^6^ uveitogenic leukocytes promotes uveitis in mice with and without AIA.**

EAU was induced in female C57BL/6 Ly5 (CD45.1^+^) mice before the recovery of splenic and lymph node leukocytes at day 11 post immunization. Prior to transfer, leukocytes were cultured for 3 days in the presence of RBP-3_629-643_ and IL-23, with IL-2 added after the first 24 h of culture. 1 x 10^6^ leukocytes were transferred into recipient C57BL/6 WT (CD45.2^+^) control mice without AIA, or mice at day 10 of AIA. **(A-B)** Frequency of EAU incidence (A) and OCT clinical disease scores (B) were determined at day 7 post cell transfer (n = 12 eyes per group). **(C)** Representative OCT imaging of control mice and mice with AIA at day 9 of AIA prior to adoptive leukocyte transfer on day 10, and at day 7 post cell transfer (day 17 post AIA induction) (R, retina; V, vitreous; C arrowhead, cellular infiltrate). **(D)** Flow cytometric quantification of total CD45^+^, transferred (CD45.1^+^) and endogenous (CD45.2^+^) ocular leukocytes at day 7 post cell transfer (n = 9-16 eyes per group). **(E)** Flow cytometric quantification of total ocular CD4^+^ T cells and CD11b^+^ myeloid cells at day 7 post cell transfer (n = 9-16 eyes per group). Graphs show mean ±SD.

## Notes

### Competing Interest Statement

The authors have declared no competing interest.

## References

[1] G. I. Collaborators, Global, regional, and national incidence of six major immune-mediated inflammatory diseases: findings from the global burden of disease study 2019, EClinicalMedicine. 64 (2023) 102193. 10.1016/j.eclinm.2023.102193

[2] N. Conrad, S. Misra, J. Y. Verbakel, G. Verbeke, G. Molenberghs, P. N. Taylor, J. Mason, N. Sattar, J. J. V. McMurray, I. B. McInnes, K. Khunti, G. Cambridge, Incidence, prevalence, and co-occurrence of autoimmune disorders over time and by age, sex, and socioeconomic status: a population-based cohort study of 22 million individuals in the UK, Lancet. 401 (2023) 1878–1890. 10.1016/S0140-6736(23)00457-9

[3] S. L. N. Clarke, P. Maghsoudlou, C. M. Guly, A. D. Dick, A. V. Ramanan, The management of adult and paediatric uveitis for rheumatologists, Nat Rev Rheumatol. 20 (2024) 795–808. 10.1038/s41584-024-01181-x

[4] E. Miserocchi, G. Fogliato, G. Modorati, F. Bandello, Review on the worldwide epidemiology of uveitis, Eur J Ophthalmol. 23 (2013) 705–717. 10.5301/ejo.5000278

[5] N. Zeboulon, M. Dougados, L. Gossec, Prevalence and characteristics of uveitis in the spondyloarthropathies: a systematic literature review, Ann Rheum Dis. 67 (2008) 955–959. 10.1136/ard.2007.075754

[6] J. Gueudry, S. Touhami, P. Quartier, B. Bodaghi, Therapeutic advances in juvenile idiopathic arthritis - associated uveitis, Curr Opin Ophthalmol. 30 (2019) 179–186. 10.1097/ICU.0000000000000559

[7] E. S. Sen, A. Ramanan, Juvenile idiopathic arthritis-associated uveitis, Clinical Immunology. 211 (2020) 108322.

[8] G. M. Leone, K. Mangano, M. C. Petralia, F. Nicoletti, P. Fagone, Past, Present and (Foreseeable) Future of Biological Anti-TNF Alpha Therapy, J Clin Med. 12 (2023) 10.3390/jcm12041630

[9] S. Siebert, A. Tsoukas, J. Robertson, I. McInnes, Cytokines as therapeutic targets in rheumatoid arthritis and other inflammatory diseases, Pharmacol Rev. 67 (2015) 280–309. 10.1124/pr.114.009639

[10] Z. Szekanecz, I. B. McInnes, G. Schett, S. Szamosi, S. Benko, G. Szucs, Autoinflammation and autoimmunity across rheumatic and musculoskeletal diseases, Nat Rev Rheumatol. 17 (2021) 585–595. 10.1038/s41584-021-00652-9

[11] E. H. Choy, F. De Benedetti, T. Takeuchi, M. Hashizume, M. R. John, T. Kishimoto, Translating IL- 6 biology into effective treatments, Nat Rev Rheumatol. 16 (2020) 335–345. 10.1038/s41584-020-0419-z

[12] M. Hassan, M. A. Sadiq, M. S. Ormaechea, G. Uludag, M. S. Halim, R. Afridi, D. V. Do, Y. J. Sepah, Q. D. Nguyen, Utilisation of composite endpoint outcome to assess efficacy of tocilizumab for non-infectious uveitis in the STOP-Uveitis Study, Br J Ophthalmol. 107 (2023) 1197–1201. 10.1136/bjophthalmol-2021-320604

[13] Y. J. Sepah, M. A. Sadiq, D. S. Chu, M. Dacey, R. Gallemore, P. Dayani, M. Hanout, M. Hassan, R. Afridi, A. Agarwal, M. S. Halim, D. V. Do, Q. D. Nguyen, Primary (Month-6) Outcomes of the STOP-Uveitis Study: Evaluating the Safety, Tolerability, and Efficacy of Tocilizumab in Patients With Noninfectious Uveitis, Am J Ophthalmol. 183 (2017) 71–80. 10.1016/j.ajo.2017.08.019

[14] D. G. McGonagle, I. B. McInnes, B. W. Kirkham, J. Sherlock, R. Moots, The role of IL-17A in axial spondyloarthritis and psoriatic arthritis: recent advances and controversies, Ann Rheum Dis. 78 (2019) 1167–1178. 10.1136/annrheumdis-2019-215356

[15] C. H. Bang, H. J. Oh, Y. H. Kim, J. H. Jung, J. H. Lee, Y. M. Park, J. H. Han, Ustekinumab Demonstrates Lower Uveitis Risk in Moderate to Severe Psoriasis Patients Compared with Tumor Necrosis Factor-alpha Inhibitors, Acta Derm Venereol. 104 (2024) adv34206. 10.2340/actadv.v104.34206

[16] M. A. Brown, M. Rudwaleit, F. A. van Gaalen, N. Haroon, L. S. Gensler, C. Fleurinck, A. Marten, U. Massow, N. de Peyrecave, T. Vaux, K. White, A. Deodhar, I. van der Horst-Bruinsma, Low uveitis rates in patients with axial spondyloarthritis treated with bimekizumab: pooled results from phase 2b/3 trials, Ann Rheum Dis. 83 (2024) 1722–1730. 10.1136/ard-2024-225933

[17] E. Letko, S. Yeh, C. S. Foster, U. Pleyer, M. Brigell, C. L. Grosskreutz, A. A. S. Group, Efficacy and safety of intravenous secukinumab in noninfectious uveitis requiring steroid-sparing immunosuppressive therapy, Ophthalmology. 122 (2015) 939–948. 10.1016/j.ophtha.2014.12.033

[18] A. Amadi-Obi, C. R. Yu, X. Liu, R. M. Mahdi, G. L. Clarke, R. B. Nussenblatt, I. Gery, Y. S. Lee, C. E. Egwuagu, TH17 cells contribute to uveitis and scleritis and are expanded by IL-2 and inhibited by IL-27/STAT1, Nat Med. 13 (2007) 711–718. 10.1038/nm1585

[19] D. Luger, P. B. Silver, J. Tang, D. Cua, Z. Chen, Y. Iwakura, E. P. Bowman, N. M. Sgambellone, C. C. Chan, R. R. Caspi, Either a Th17 or a Th1 effector response can drive autoimmunity: conditions of disease induction affect dominant effector category, J Exp Med. 205 (2008) 799–810. 10.1084/jem.20071258

[20] D. Sun, D. Liang, H. J. Kaplan, H. Shao, The role of Th17-associated cytokines in the pathogenesis of experimental autoimmune uveitis (EAU), Cytokine. 74 (2015) 76–80. 10.1016/j.cyto.2014.12.017

[21] K. Bechman, Z. Yang, M. Adas, D. Nagra, S. U. A, M. D. Russell, N. Wilson, S. Steer, S. Norton, J. Galloway, Incidence of Uveitis in Patients With Axial Spondylarthritis Treated With Biologics or Targeted Synthetics: A Systematic Review and Network Meta-Analysis, Arthritis Rheumatol. 76 (2024) 704–714. 10.1002/art.42788

[22] P. Quartier, A. Baptiste, V. Despert, E. Allain-Launay, I. Kone-Paut, A. Belot, L. Kodjikian, D. Monnet, M. Weber, C. Elie, B. Bodaghi, A. S. Group, ADJUVITE: a double-blind, randomised, placebo-controlled trial of adalimumab in early onset, chronic, juvenile idiopathic arthritis-associated anterior uveitis, Ann Rheum Dis. 77 (2018) 1003–1011. 10.1136/annrheumdis-2017-212089

[23] A. V. Ramanan, A. D. Dick, C. Guly, A. McKay, A. P. Jones, B. Hardwick, R. W. J. Lee, M. Smyth, T. Jaki, M. W. Beresford, A. T. M. Group, Tocilizumab in patients with anti-TNF refractory juvenile idiopathic arthritis-associated uveitis (APTITUDE): a multicentre, single-arm, phase 2 trial, Lancet Rheumatol. 2 (2020) e135–e141. 10.1016/S2665-9913(20)30008-4

[24] C. Tappeiner, M. Mesquida, A. Adan, J. Anton, A. V. Ramanan, E. Carreno, F. Mackensen, K. Kotaniemi, J. H. de Boer, R. Bou, C. G. de Vicuna, A. Heiligenhaus, Evidence for Tocilizumab as a Treatment Option in Refractory Uveitis Associated with Juvenile Idiopathic Arthritis, J Rheumatol. 43 (2016) 2183–2188. 10.3899/jrheum.160231

[25] E. J. Lee, E. E. Vance, B. R. Brown, P. S. Snow, J. S. Clowers, S. Sakaguchi, H. L. Rosenzweig, Investigation of the relationship between the onset of arthritis and uveitis in genetically predisposed SKG mice, Arthritis Res Ther. 17 (2015) 218. 10.1186/s13075-015-0725-z

[26] S. C. Osinchuk, B. H. Grahn, T. D. Wilson, B. N. Thompson, D. A. Hart, K. D. Harrison, D. M. Cooper, A. Panahifar, A. M. Rosenberg, Evaluation of Uveitis Induced in Rats by a Type I Collagen Peptide as a Model for Childhood Arthritis-associated Uveitis, Comp Med. 73 (2023) 267–276. 10.30802/AALAS-CM-22-000129

[27] H. L. Rosenzweig, T. M. Martin, S. R. Planck, M. M. Jann, J. R. Smith, T. T. Glant, W. van Eden, M. P. Davey, J. T. Rosenbaum, Anterior uveitis accompanies joint disease in a murine model resembling ankylosing spondylitis, Ophthalmic Res. 40 (2008) 189–192. 10.1159/000119874

[28] M. Ruutu, G. Thomas, R. Steck, M. A. Degli-Esposti, M. S. Zinkernagel, K. Alexander, J. Velasco, G. Strutton, A. Tran, H. Benham, L. Rehaume, R. J. Wilson, K. Kikly, J. Davies, A. R. Pettit, M. A. Brown, M. A. McGuckin, R. Thomas, beta-glucan triggers spondylarthritis and Crohn’s disease-like ileitis in SKG mice, Arthritis Rheum. 64 (2012) 2211–2222. 10.1002/art.34423

[29] G. W. Jones, M. Bombardieri, C. J. Greenhill, L. McLeod, A. Nerviani, V. Rocher-Ros, A. Cardus, A. S. Williams, C. Pitzalis, B. J. Jenkins, S. A. Jones, Interleukin-27 inhibits ectopic lymphoid-like structure development in early inflammatory arthritis, J Exp Med. 212 (2015) 1793–1802. 10.1084/jem.20132307

[30] M. A. Nowell, A. S. Williams, S. A. Carty, J. Scheller, A. J. Hayes, G. W. Jones, P. J. Richards, S. Slinn, M. Ernst, B. J. Jenkins, N. Topley, S. Rose-John, S. A. Jones, Therapeutic targeting of IL-6 trans signaling counteracts STAT3 control of experimental inflammatory arthritis, J Immunol. 182 (2009) 613–622. 10.4049/jimmunol.182.1.613

[31] S. Ohshima, Y. Saeki, T. Mima, M. Sasai, K. Nishioka, S. Nomura, M. Kopf, Y. Katada, T. Tanaka, M. Suemura, T. Kishimoto, Interleukin 6 plays a key role in the development of antigen-induced arthritis, Proc Natl Acad Sci U S A. 95 (1998) 8222–8226. 10.1073/pnas.95.14.8222

[32] S. R. Pickens, N. D. Chamberlain, M. V. Volin, A. M. Mandelin, 2nd, H. Agrawal, M. Matsui, T. Yoshimoto, S. Shahrara, Local expression of interleukin-27 ameliorates collagen-induced arthritis, Arthritis Rheum. 63 (2011) 2289–2298. 10.1002/art.30324

[33] J. P. Twohig, A. Cardus Figueras, R. Andrews, F. Wiede, B. C. Cossins, A. Derrac Soria, M. J. Lewis, M. J. Townsend, D. Millrine, J. Li, D. G. Hill, J. Uceda Fernandez, X. Liu, B. Szomolay, C. J. Pepper, P. R. Taylor, C. Pitzalis, T. Tiganis, N. M. Williams, G. W. Jones, S. A. Jones, Activation of naive CD4(+) T cells re-tunes STAT1 signaling to deliver unique cytokine responses in memory CD4(+) T cells, Nat Immunol. 20 (2019) 458–470. 10.1038/s41590-019-0350-0

[34] G. W. Jones, R. M. McLoughlin, V. J. Hammond, C. R. Parker, J. D. Williams, R. Malhotra, J. Scheller, A. S. Williams, S. Rose-John, N. Topley, S. A. Jones, Loss of CD4+ T cell IL-6R expression during inflammation underlines a role for IL-6 trans signaling in the local maintenance of Th17 cells, J Immunol. 184 (2010) 2130–2139. 10.4049/jimmunol.0901528

[35] H. Yoshida, S. Hamano, G. Senaldi, T. Covey, R. Faggioni, S. Mu, M. Xia, A. C. Wakeham, H. Nishina, J. Potter, C. J. Saris, T. W. Mak, WSX-1 is required for the initiation of Th1 responses and resistance to L. major infection, Immunity. 15 (2001) 569–578. 10.1016/s1074-7613(01)00206-0

[36] O. H. Bell, D. A. Copland, A. Ward, L. B. Nicholson, C. A. K. Lange, C. J. Chu, A. D. Dick, Single Eye mRNA-Seq Reveals Normalisation of the Retinal Microglial Transcriptome Following Acute Inflammation, Front Immunol. 10 (2019) 3033. 10.3389/fimmu.2019.03033

[37] G. W. Jones, D. G. Hill, K. Sime, A. S. Williams, In Vivo Models for Inflammatory Arthritis, Methods Mol Biol. 1725 (2018) 101–118. 10.1007/978-1-4939-7568-6_9

[38] J. Boldison, T. K. Khera, D. A. Copland, M. L. Stimpson, G. L. Crawford, A. D. Dick, L. B. Nicholson, A novel pathogenic RBP-3 peptide reveals epitope spreading in persistent experimental autoimmune uveoretinitis, Immunology. 146 (2015) 301–311. 10.1111/imm.12503

[39] A. Ward, O. H. Bell, L. Martinez-Robles, P. J. P. Lait, C. J. Chu, D. A. Copland, A. D. Dick, L. B. Nicholson, A pathogenic CD4 T cell phenotype in experimental uveitis shares common features with other immune mediated inflammatory diseases, Discov Immunol. 5 (2026) kyaf019. 10.1093/discim/kyaf019

[40] A. Shome, O. O. Mugisho, R. L. Niederer, I. D. Rupenthal, Comprehensive Grading System for Experimental Autoimmune Uveitis in Mice, Biomedicines. 11 (2023) 10.3390/biomedicines11072022

[41] E. C. Kerr, B. J. Raveney, D. A. Copland, A. D. Dick, L. B. Nicholson, Analysis of retinal cellular infiltrate in experimental autoimmune uveoretinitis reveals multiple regulatory cell populations, J Autoimmun. 31 (2008) 354–361. 10.1016/j.jaut.2008.08.006

[42] E. J. Liebling, W. Faig, J. C. Chang, E. Mendoza, N. Moore, N. L. Vicioso, M. A. Lerman, Temporal Relationship Between Juvenile Idiopathic Arthritis Disease Activity and Uveitis Disease Activity, Arthritis Care Res (Hoboken). 74 (2022) 349–354. 10.1002/acr.24483

[43] M. A. Nowell, P. J. Richards, S. Horiuchi, N. Yamamoto, S. Rose-John, N. Topley, A. S. Williams, S. A. Jones, Soluble IL-6 receptor governs IL-6 activity in experimental arthritis: blockade of arthritis severity by soluble glycoprotein 130, J Immunol. 171 (2003) 3202–3209. 10.4049/jimmunol.171.6.3202

[44] H. Haruta, N. Ohguro, M. Fujimoto, S. Hohki, F. Terabe, S. Serada, S. Nomura, K. Nishida, T. Kishimoto, T. Naka, Blockade of interleukin-6 signaling suppresses not only th17 but also interphotoreceptor retinoid binding protein-specific Th1 by promoting regulatory T cells in experimental autoimmune uveoretinitis, Invest Ophthalmol Vis Sci. 52 (2011) 3264–3271. 10.1167/iovs.10-6272

[45] S. Wu, R. Ma, Y. Zhong, Z. Chen, H. Zhou, M. Zhou, W. Chong, J. Chen, Deficiency of IL-27 Signaling Exacerbates Experimental Autoimmune Uveitis with Elevated Uveitogenic Th1 and Th17 Responses, Int J Mol Sci. 22 (2021) 10.3390/ijms22147517

[46] C. J. Chu, P. Herrmann, L. S. Carvalho, S. E. Liyanage, J. W. Bainbridge, R. R. Ali, A. D. Dick, U. F. Luhmann, Assessment and in vivo scoring of murine experimental autoimmune uveoretinitis using optical coherence tomography, PLoS One. 8 (2013) e63002. 10.1371/journal.pone.0063002

[47] R. R. Caspi, A look at autoimmunity and inflammation in the eye, J Clin Invest. 120 (2010) 3073–3083. 10.1172/JCI42440

[48] J. Cunha-Vaz, R. Bernardes, C. Lobo, Blood-retinal barrier, Eur J Ophthalmol. 21 Suppl 6 (2011) S3-9. 10.5301/EJO.2010.6049

[49] K. Baquet-Walscheid, K. Minden, M. Niewerth, F. Dressler, I. Foeldvari, D. Foell, J. P. Haas, G. Horneff, A. Hospach, T. Kallinich, J. Kummerle-Deschner, K. Monkemoller, C. Tappeiner, D. Windschall, J. Klotsche, A. Heiligenhaus, Course of uveitis in children with juvenile idiopathic arthritis (JIA): Five years follow-up data from a prospective multicenter Inception Cohort of Newly diagnosed patients with JIA (ICON-JIA) study, Arthritis Res Ther. 27 (2025) 61. 10.1186/s13075-025-03531-w

[50] L. Li, J. M. Zhang, J. H. Deng, W. Y. Kuang, X. H. Tan, C. Li, S. P. Li, C. F. Li, Clinical Characteristics and Prognosis of Juvenile Idiopathic Arthritis-Associated Uveitis: A Single-Center Retrospective Study, Clin Ophthalmol. 19 (2025) 2813–2820. 10.2147/OPTH.S529421

[51] M. E. Zannin, I. Buscain, F. Vittadello, G. Martini, M. Alessio, J. G. Orsoni, L. Breda, D. Rigante, R. Cimaz, F. Zulian, Timing of uveitis onset in oligoarticular juvenile idiopathic arthritis (JIA) is the main predictor of severe course uveitis, Acta Ophthalmol. 90 (2012) 91–95. 10.1111/j.1755-3768.2009.01815.x

[52] N. Sakaguchi, T. Takahashi, H. Hata, T. Nomura, T. Tagami, S. Yamazaki, T. Sakihama, T. Matsutani, I. Negishi, S. Nakatsuru, S. Sakaguchi, Altered thymic T-cell selection due to a mutation of the ZAP-70 gene causes autoimmune arthritis in mice, Nature. 426 (2003) 454–460. 10.1038/nature02119

[53] S. T. Angeles-Han, V. M. Utz, S. Thornton, G. Schulert, J. Rodriguez-Smith, A. Kauffman, A. Sproles, N. Mwase, T. Hennard, A. Grom, M. Altaye, G. N. Holland, S100 proteins, cytokines, and chemokines as tear biomarkers in children with juvenile idiopathic arthritis-associated uveitis, Ocul Immunol Inflamm. 29 (2021) 1616–1620. 10.1080/09273948.2020.1758731

[54] C. Tappeiner, J. Klotsche, C. Sengler, M. Niewerth, I. Liedmann, K. Walscheid, M. Lavric, D. Foell, K. Minden, A. Heiligenhaus, Risk Factors and Biomarkers for the Occurrence of Uveitis in Juvenile Idiopathic Arthritis: Data From the Inception Cohort of Newly Diagnosed Patients With Juvenile Idiopathic Arthritis Study, Arthritis Rheumatol. 70 (2018) 1685–1694. 10.1002/art.40544

[55] K. Walscheid, A. Heiligenhaus, D. Holzinger, J. Roth, C. Heinz, C. Tappeiner, M. Kasper, D. Foell, Elevated S100A8/A9 and S100A12 Serum Levels Reflect Intraocular Inflammation in Juvenile Idiopathic Arthritis-Associated Uveitis: Results From a Pilot Study, Invest Ophthalmol Vis Sci. 56 (2015) 7653–7660. 10.1167/iovs.15-17066

[56] O. C. Kwon, E. J. Lee, J. Y. Lee, J. Youn, T. H. Kim, S. Hong, C. K. Lee, B. Yoo, W. H. Robinson, Y. G. Kim, Prefoldin 5 and Anti-prefoldin 5 Antibodies as Biomarkers for Uveitis in Ankylosing Spondylitis, Front Immunol. 10 (2019) 384. 10.3389/fimmu.2019.00384

[57] P. Di Meglio, A. Di Cesare, U. Laggner, C. C. Chu, L. Napolitano, F. Villanova, I. Tosi, F. Capon, R. C. Trembath, K. Peris, F. O. Nestle, The IL23R R381Q gene variant protects against immune-mediated diseases by impairing IL-23-induced Th17 effector response in humans, PLoS One. 6 (2011) e17160. 10.1371/journal.pone.0017160

[58] R. Sarin, X. Wu, C. Abraham, Inflammatory disease protective R381Q IL23 receptor polymorphism results in decreased primary CD4+ and CD8+ human T-cell functional responses, Proc Natl Acad Sci U S A. 108 (2011) 9560–9565. 10.1073/pnas.1017854108

[59] P. K. Wong, J. M. Quinn, N. A. Sims, A. van Nieuwenhuijze, I. K. Campbell, I. P. Wicks, Interleukin-6 modulates production of T lymphocyte-derived cytokines in antigen-induced arthritis and drives inflammation-induced osteoclastogenesis, Arthritis Rheum. 54 (2006) 158–168. 10.1002/art.21537

[60] W. Razawy, P. S. Asmawidjaja, A. M. Mus, N. Salioska, N. Davelaar, N. Kops, M. Oukka, C. H. Alves, E. Lubberts, CD4(+) CCR6(+) T cells, but not gammadelta T cells, are important for the IL-23R-dependent progression of antigen-induced inflammatory arthritis in mice, Eur J Immunol. 50 (2020) 245–255. 10.1002/eji.201948112

[61] M. I. Koenders, E. Lubberts, B. Oppers-Walgreen, L. van den Bersselaar, M. M. Helsen, F. E. Di Padova, A. M. Boots, H. Gram, L. A. Joosten, W. B. van den Berg, Blocking of interleukin-17 during reactivation of experimental arthritis prevents joint inflammation and bone erosion by decreasing RANKL and interleukin-1, Am J Pathol. 167 (2005) 141–149. 10.1016/S0002-9440(10)62961-6

[62] K. Hirahara, A. Onodera, A. V. Villarino, M. Bonelli, G. Sciume, A. Laurence, H. W. Sun, S. R. Brooks, G. Vahedi, H. Y. Shih, G. Gutierrez-Cruz, S. Iwata, R. Suzuki, Y. Mikami, Y. Okamoto, T. Nakayama, S. M. Holland, C. A. Hunter, Y. Kanno, J. J. O’Shea, Asymmetric Action of STAT Transcription Factors Drives Transcriptional Outputs and Cytokine Specificity, Immunity. 42 (2015) 877–889. 10.1016/j.immuni.2015.04.014

[63] S. A. Jones, B. J. Jenkins, Recent insights into targeting the IL-6 cytokine family in inflammatory diseases and cancer, Nat Rev Immunol. 18 (2018) 773–789. 10.1038/s41577-018-0066-7

[64] A. V. Villarino, Y. Kanno, J. J. O’Shea, Mechanisms and consequences of Jak-STAT signaling in the immune system, Nat Immunol. 18 (2017) 374–384. 10.1038/ni.3691

[65] J. S. Stumhofer, A. Laurence, E. H. Wilson, E. Huang, C. M. Tato, L. M. Johnson, A. V. Villarino, Q. Huang, A. Yoshimura, D. Sehy, C. J. Saris, J. J. O’Shea, L. Hennighausen, M. Ernst, C. A. Hunter, Interleukin 27 negatively regulates the development of interleukin 17-producing T helper cells during chronic inflammation of the central nervous system, Nat Immunol. 7 (2006) 937–945. 10.1038/ni1376

[66] M. Mesquida, F. Drawnel, P. J. Lait, D. A. Copland, M. L. Stimpson, V. Llorenc, M. Sainz de la Maza, A. Adan, G. Widmer, P. Strassburger, S. Fauser, A. D. Dick, R. W. J. Lee, B. Molins, Modelling Macular Edema: The Effect of IL-6 and IL-6R Blockade on Human Blood-Retinal Barrier Integrity In Vitro, Transl Vis Sci Technol. 8 (2019) 32. 10.1167/tvst.8.5.32

[67] S. Sharma, Interleukin-6 Trans-signaling: A Pathway With Therapeutic Potential for Diabetic Retinopathy, Front Physiol. 12 (2021) 689429. 10.3389/fphys.2021.689429

[68] M. L. Valle, J. Dworshak, A. Sharma, A. S. Ibrahim, M. Al-Shabrawey, S. Sharma, Inhibition of interleukin-6 trans-signaling prevents inflammation and endothelial barrier disruption in retinal endothelial cells, Exp Eye Res. 178 (2019) 27–36. 10.1016/j.exer.2018.09.009

[69] S. Sharma, P. Lin, A. Hu, E. B. Suhler, M. Pauly-Evers, W. Holmes, Z. Barekati, D. Willen, L. Macgregor, L. Steeples, Z. Haskova, D. Feenstra, D. Silverman, B. Passemard, S. Fauser, M. Mesquida, Interleukin 6 Inhibition With Vamikibart for Uveitic Macular Edema: The Phase 1 DOVETAIL Nonrandomized Clinical Trial, JAMA Ophthalmol. 144 (2026) 442–451. 10.1001/jamaophthalmol.2026.0610

