## Supplementary Data for "Inflammatory arthritis disrupts ocular immune privilege by compromising blood-retinal barrier integrity and promoting uveitogenic T cell recruitment"

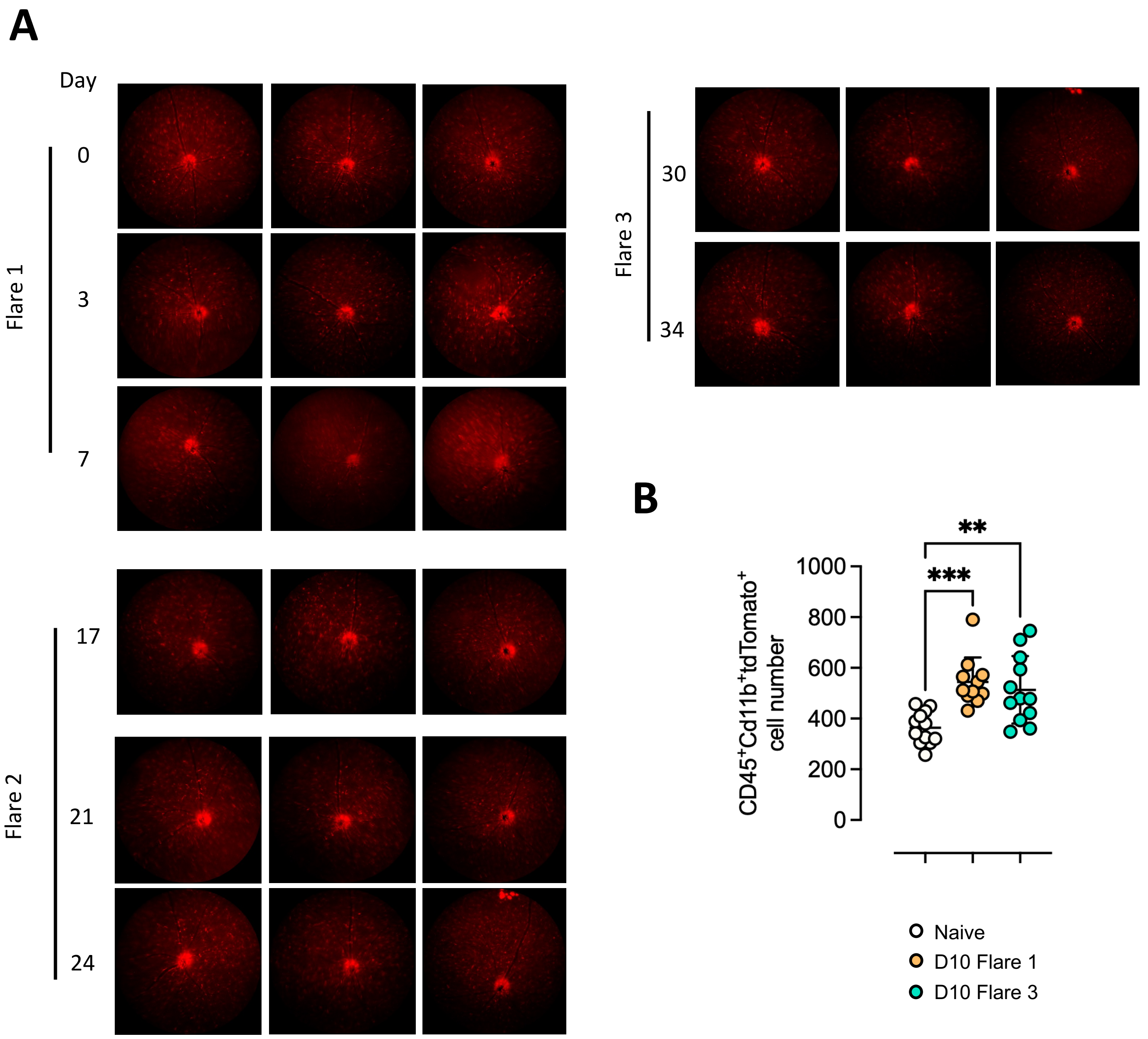


***Supplemental Figure 1.*** *Resident ocular microglia numbers increase after AIA induction.*

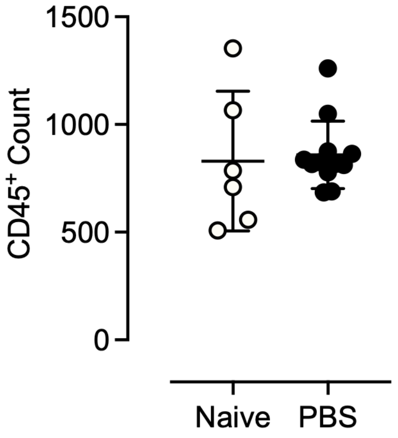


***Supplemental Figure 2.*** *Intra-articular administration of PBS does not promote ocular leukocyte recruitment*

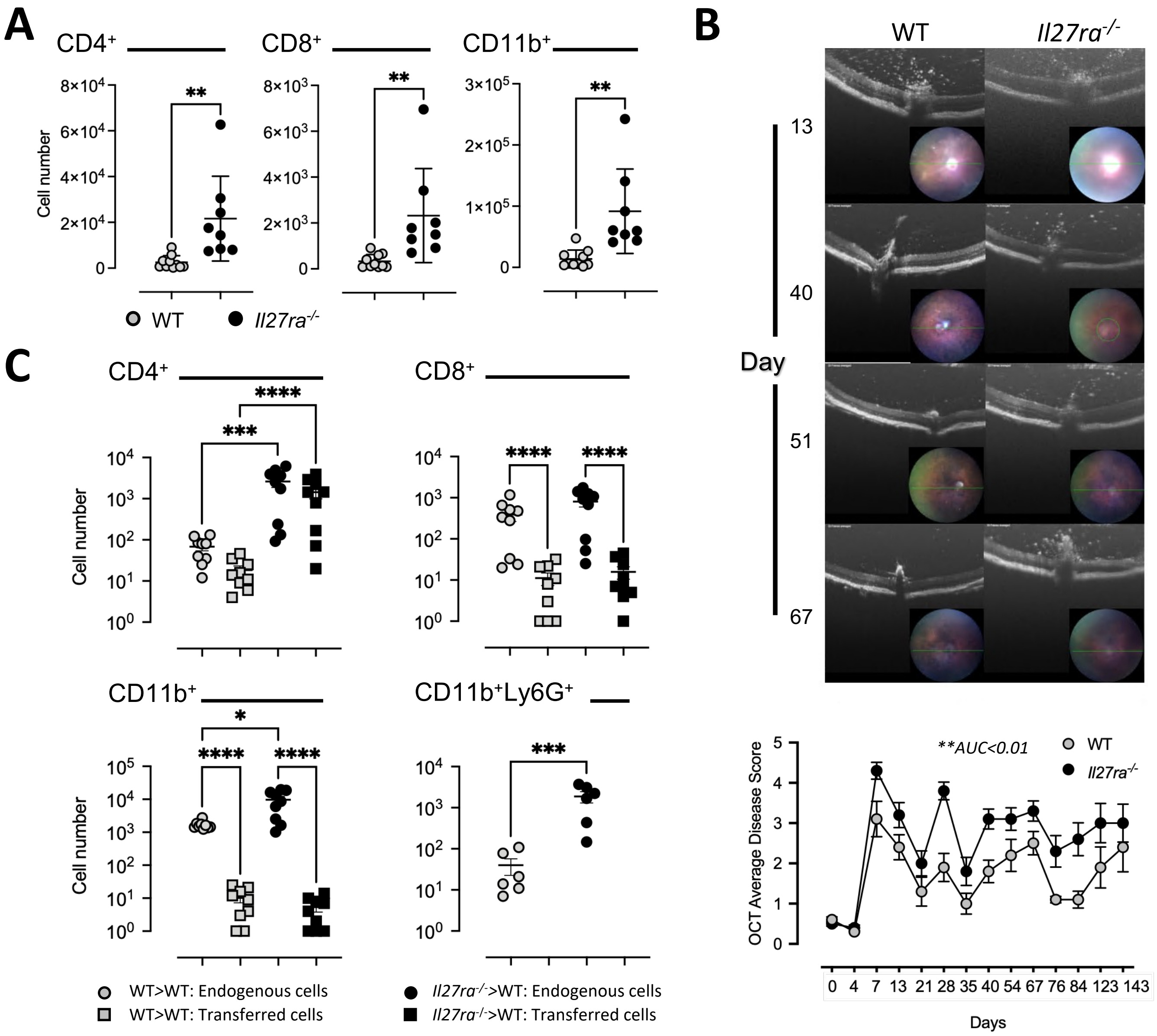


***Supplemental Figure 3.*** *IL-27 signaling limits the severity of EAU.*

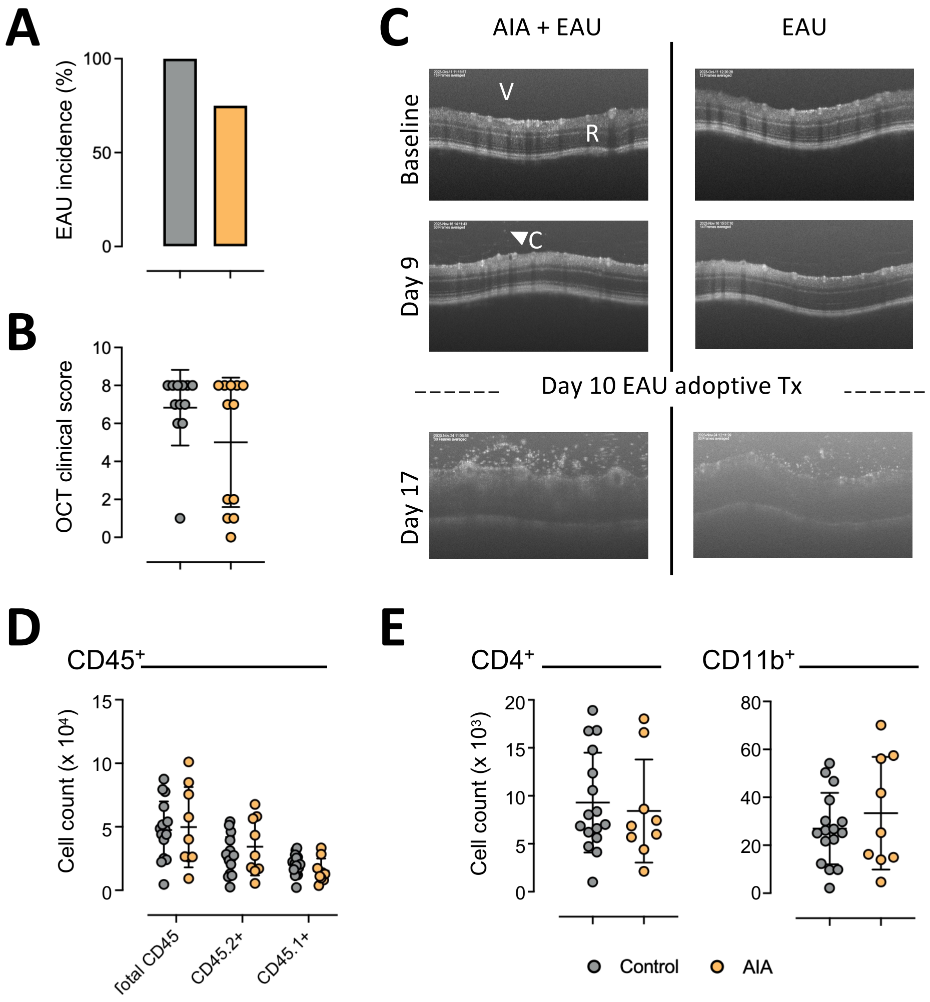


***Supplemental Figure 4.*** *Adoptive transfer of 1 x 10$^{6}$ uveitogenic leukocytes promotes uveitis in mice with and without AIA.*

| Target | Conjugate | Clone | Supplier | Catalogue Number |
|---|---|---|---|---|
| CD45 | FITC | 30-F11 | Invitrogen | 11-0451-82 |
| CD3 | PE-Cy7 | 145-2C11 | eBioscience | MA5-17655 |
| CD4 | APC | RM4-5 | Invitrogen | MCD0405 |
| CD8 | AF700 | 53-6.7 | R&D Systems | 100729 |
| LY6G | V450 | 1A8 | BD | 560603 |
| CD11B | APC-Cy7 | M1/70 | BD | 055657 |
| 7AAD | N/A | N/A | Biolegend | 420404 |
| CD45.1 | BV605 | A20 | Biolegend | 110738 |

**Supplemental Table 1**

**Fluorochrome-conjugated antibodies used to characterize ocular leukocytes in AIA**
